# Biosynthetic gene clusters, secondary metabolite profiles, and cards of virulence in the closest nonpathogenic relatives of *Aspergillus fumigatus*

**DOI:** 10.1101/2020.04.09.033902

**Authors:** Jacob L. Steenwyk, Matthew E. Mead, Sonja L. Knowles, Huzefa A. Raja, Christopher D. Roberts, Oliver Bader, Jos Houbraken, Gustavo H. Goldman, Nicholas H. Oberlies, Antonis Rokas

## Abstract

*Aspergillus fumigatus* is a major human pathogen that causes hundreds of thousands of infections yearly with high mortality rates. In contrast, *Aspergillus fischeri* and the recently described *Aspergillus oerlinghausenensis*, the two species most closely related to *A. fumigatus*, are not known to be pathogenic. Some of the “cards of virulence” that A. *fumigatus* possesses are secondary metabolites that impair the host immune system, protect from host immune cell attacks, or acquire key nutrients. Secondary metabolites and the biosynthetic gene clusters (BGCs) that typically encode them often vary within and between fungal species. To gain insight into whether secondary metabolism-associated cards of virulence vary between *A. fumigatus, A. oerlinghausenensis*, and *A. fischeri*, we conducted extensive genomic and secondary metabolite profiling analyses. By analyzing multiple *A. fumigatus*, one *A. oerlinghausenensis*, and multiple *A. fischeri* strains, we identified both conserved and diverged secondary metabolism-associated cards of virulence. For example, we found that all species and strains examined biosynthesized the major virulence factor gliotoxin, consistent with the conservation of the gliotoxin BGC across genomes. However, species differed in their biosynthesis of fumagillin and pseurotin, both contributors to host tissue damage during invasive aspergillosis; these differences were reflected in sequence divergence of the intertwined fumagillin/pseurotin BGCs across genomes. These results delineate the similarities and differences in secondary metabolism-associated cards of virulence between a major fungal pathogen and its nonpathogenic closest relatives, shedding light into the genetic and phenotypic changes associated with the evolution of fungal pathogenicity.

**Importance:** The major fungal pathogen *Aspergillus fumigatus* kills tens of thousands each year. In contrast, the two closest relatives of *A. fumigatus*, namely *Aspergillus fischeri* and *Aspergillus oerlinghausenensis*, are not considered pathogenic. *A. fumigatus* virulence stems, partly, from its ability to produce small molecules called secondary metabolites that have potent activities during infection. In this study, we examined whether *A. fumigatus* secondary metabolites and the metabolic pathways involved in their production are conserved in *A. oerlinghausenensis* and *A. fischeri*. We found that the nonpathogenic close relatives of *A. fumigatus* produce some, but not all, secondary metabolites thought to contribute to the success of *A. fumigatus* in causing human disease and that these similarities and differences were reflected in the underlying metabolic pathways involved in their biosynthesis. Compared to its nonpathogenic close relatives, *A. fumigatus* produces a distinct cocktail of secondary metabolites, which likely contributes to these organisms’ vastly different potentials to cause human disease. More broadly, the study of nonpathogenic organisms that have virulence-related traits, but are not currently considered agents of human disease, may facilitate the prediction of species capable of posing future threats to human health.

## Introduction

Fungal diseases impose a clinical, economic, and social burden on humans (Drgona et al., 2014; Vallabhaneni et al., 2016; Benedict et al., 2019). Fungi from the genus *Aspergillus* are responsible for a considerable fraction of this burden, accounting for more than 250,000 infections annually with high mortality rates (Bongomin et al., 2017). *Aspergillus* infections often result in pulmonary and invasive diseases that are collectively termed aspergillosis. Among *Aspergillus* species, *Aspergillus fumigatus* is the primary etiological agent of aspergillosis (Latgé and Chamilos, 2019).

Even though *A. fumigatus* is a major pathogen, its closest relatives are not considered pathogenic (Mead et al., 2019a; Steenwyk et al., 2019; Rokas et al., 2020a). Numerous studies have identified factors that contribute to *A. fumigatus* pathogenicity, such as the organism’s ability to grow well at higher temperatures and in hypoxic conditions (Kamei and Watanabe, 2005; Tekaia and Latgé, 2005; Abad et al., 2010; Grahl et al., 2012). Factors that contribute to pathogenicity could be conceived as analogous to individual “cards” of a “hand” (set of cards) in a card game – that is, individual factors are typically insufficient to cause disease but can collectively do so (Casadevall, 2007).

Several secondary metabolites biosynthesized by *A. fumigatus* are known “cards” of virulence because of their involvement in impairing the host immune system, protecting the fungus from host immune cell attacks, or acquiring key nutrients (Raffa and Keller, 2019). For example, the secondary metabolite gliotoxin has been shown to contribute to *A. fumigatus* virulence by inhibiting the host immune response (Sugui et al., 2007). Other secondary metabolites implicated in virulence include: fumitremorgin, which inhibits the activity of the breast cancer resistance protein (González-Lobato et al., 2010); verruculogen, which modulates the electrophysical properties of human nasal epithelial cells (Khoufache et al., 2007); trypacidin, which is cytotoxic to lung cells (Gauthier et al., 2012); pseurotin, which inhibits immunoglobulin E (Ishikawa et al., 2009); and fumagillin which causes epithelial cell damage (Guruceaga et al., 2018) and impairs the function of neutrophils (Fallon et al., 2010, 2011) (Table 1).

**Table 1.** Select *A. fumigatus* secondary metabolites implicated in human disease.

|  | Function | Reference(s) | Evidence of secondary metabolite |  |  |  |  |  |  |
| --- | --- | --- | --- | --- | --- | --- | --- | --- | --- |
|  |  |  | *A. fumigatus* |  |  | *A. oerlinghausensis* | *A. fischeri* |  |  |
|  |  |  | Af293 | CEA10 | CEA17 | CBS 139183<sup>T</sup> | NRRL 181 | NRRL 4585 | NRRL 4161 |
| **Gliotoxin** | Inhibits host immune response | (Sugui et al., 2007) | + | + | + | + | + | + | + |
| **Fumitremorgin** | Inhibits the breast cancer resistance protein | (González-Lobato et al., 2010) | - | + | - | + | + | + | + |
| **Verruculogen** | Changes electrophysical properties of human nasal epithelial cells | (Khoufache et al., 2007) | - | - | - | + | + | + | + |
| **Trypacidin** | Damages lung cell tissues | (Gauthier et al., 2012) | + | + | - | + | - | - | - |
| **Pseurotin** | Inhibits immunoglobulin E | (Ishikawa et al., 2009) | + | + | + | + | - | - | - |
| **Fumagillin** | Inhibits neutrophil function | (Fallon et al., 2010, 2011) | + | + | + | - | - | - | - |

A list of select secondary metabolites implicated in human disease and their functional role are described here. All secondary metabolites listed or analogs thereof were identified during secondary metabolite profiling. ‘+’ (orange) and ‘-’ (blue) indicate if the secondary metabolite was or was not produced by a strain of *A. fumigatus* or its two closest relatives.

By extension, the metabolic pathways responsible for the biosynthesis of secondary metabolites are also “cards” of virulence. Genes in these pathways are typically organized in contiguous sets termed biosynthetic gene clusters (BGCs) (Keller, 2019). BGCs are known to evolve rapidly, and their composition can differ substantially across species and strains (Lind et al., 2015, 2017; Rokas et al., 2018, 2020b). For example, even though *A. fumigatus* contains 33 BGCs and *A. fischeri* contains 48 BGCs, only 10 of those BGCs appear to be shared between the two species (Mead et al., 2019a). Interestingly, one of the BGCs that is conserved between *A. fumigatus* and *A. fischeri* is the gliotoxin BGC and both species have been shown to biosynthesize the toxic virulence factor (Knowles et al., 2020). These results suggest that the gliotoxin “card” is part of a winning “hand” that facilitates virulence only in the background of the major pathogen *A. fumigatus* and not in that of the nonpathogen *A. fischeri* (Knowles et al., 2020).

To date, such comparisons of BGCs and secondary metabolite profiles among *A. fumigatus* and closely related nonpathogenic species have been few and restricted to single strains (Mead et al., 2019a; Knowles et al., 2020). However, genetic and phenotypic heterogeneity among strains of a single species has been shown be an important factor when studying *Aspergillus* pathogenicity (Kowalski et al., 2016, 2019; Keller, 2017; Ries et al., 2019; Bastos et al., 2020; Santos et al., 2020). Examination of multiple strains of *A. fumigatus* and close relatives—including the recently described and largely uncharacterized (in the context of pathogenicity) closest known relative of *A. fumigatus, A. oerlinghausenensis* (Houbraken et al., 2016)*—*will increase our understanding of the *A. fumigatus* secondary metabolite “cards” of virulence.

To gain insight into the genomic and chemical similarities and differences in secondary metabolism among *A. fumigatus* and nonpathogenic close relatives, we characterized variation in BGCs and secondary metabolites produced by *A. fumigatus* and nonpathogenic close relatives. To do so, we first sequenced and assembled *A. oerlinghausenensis* CBS 139183^T^ as well as *A. fischeri* strains NRRL 4585 and NRRL 4161 and analyzed them together with four *A. fumigatus* and three additional *A. fischeri* publicly available genomes. We also characterized the secondary metabolite profiles of three *A. fumigatus*, one *A. oerlinghausenensis*, and three *A. fischeri* strains. On the one hand, we found that the biosynthesis of the secondary metabolites gliotoxin and fumitremorgin, which are both known to interact with mammalian cells (Yamada et al., 2000; González-Lobato et al., 2010; Li et al., 2012; Raffa and Keller, 2019), as well as their BGCs, were conserved among pathogenic and nonpathogenic strains. Interestingly, we found only *A. fischeri* strains, but not *A. fumigatus* strains, biosynthesized verruculogen, which changes the electrophysical properties of human nasal epithelial cells (Khoufache et al., 2007). On the other hand, we found that both *A. fumigatus* and *A. oerlinghausenensis* biosynthesized fumagillin and trypacidin, whose effects include broad suppression of the immune response system and lung cell damage (Ishikawa et al., 2009; Fallon et al., 2010, 2011; Gauthier et al., 2012), but *A. fischeri* did not. These results reveal that nonpathogenic close relatives of *A. fumigatus* also produce some, but not all, of the secondary metabolism-associated cards of virulence known in *A. fumigatus*. Further investigation of the similarities and differences among *A. fumigatus* and close nonpathogenic relatives may provide additional insight into the “hand of cards” that enabled *A. fumigatus* to evolve into a deadly pathogen.

## Results

### Conservation and diversity of biosynthetic gene clusters within and between species

We sequenced and assembled *A. oerlinghausenensis* CBS 139183^T^ and *A. fischeri* strains NRRL 4585 and NRRL 4161. Together with publicly available genomes, we analyzed 10 *Aspergillus* genomes (five *A. fischeri* strains; four *A. fumigatus* strains; one *A. oerlinghausenensis* strain; see Methods). We found that the newly added genomes were of similar quality to other publicly available draft genomes (average percent presence of BUSCO genes: 98.80 ± 0.10%; average N50: 451,294.67 ± 9,696.11; Fig. S1). We predicted that *A. oerlinghausenensis* CBS 139183^T^, *A. fischeri* NRRL 4585, and *A. fischeri* NRRL 4161 have 10,044, 11,152 and 10,940 genes, respectively, numbers similar to publicly available genomes. Lastly, we inferred the evolutionary history of the 10 *Aspergillus* genomes using a concatenated matrix of 3,041 genes (5,602,272 sites) and recapitulated species-level relationships as previously reported (Houbraken et al., 2016). Relaxed molecular clock analyses suggested that *A. oerlinghausenensis* CBS 139183^T^ diverged from *A. fumigatus* approximately 3.9 (6.4 – 1.3) million years ago and that *A. oerlinghausenensis* and *A. fumigatus* split from *A. fischeri* approximately 4.5 (6.8 – 1.7) million years ago (Fig. 1A; Fig. S2).

**Figure 1.**
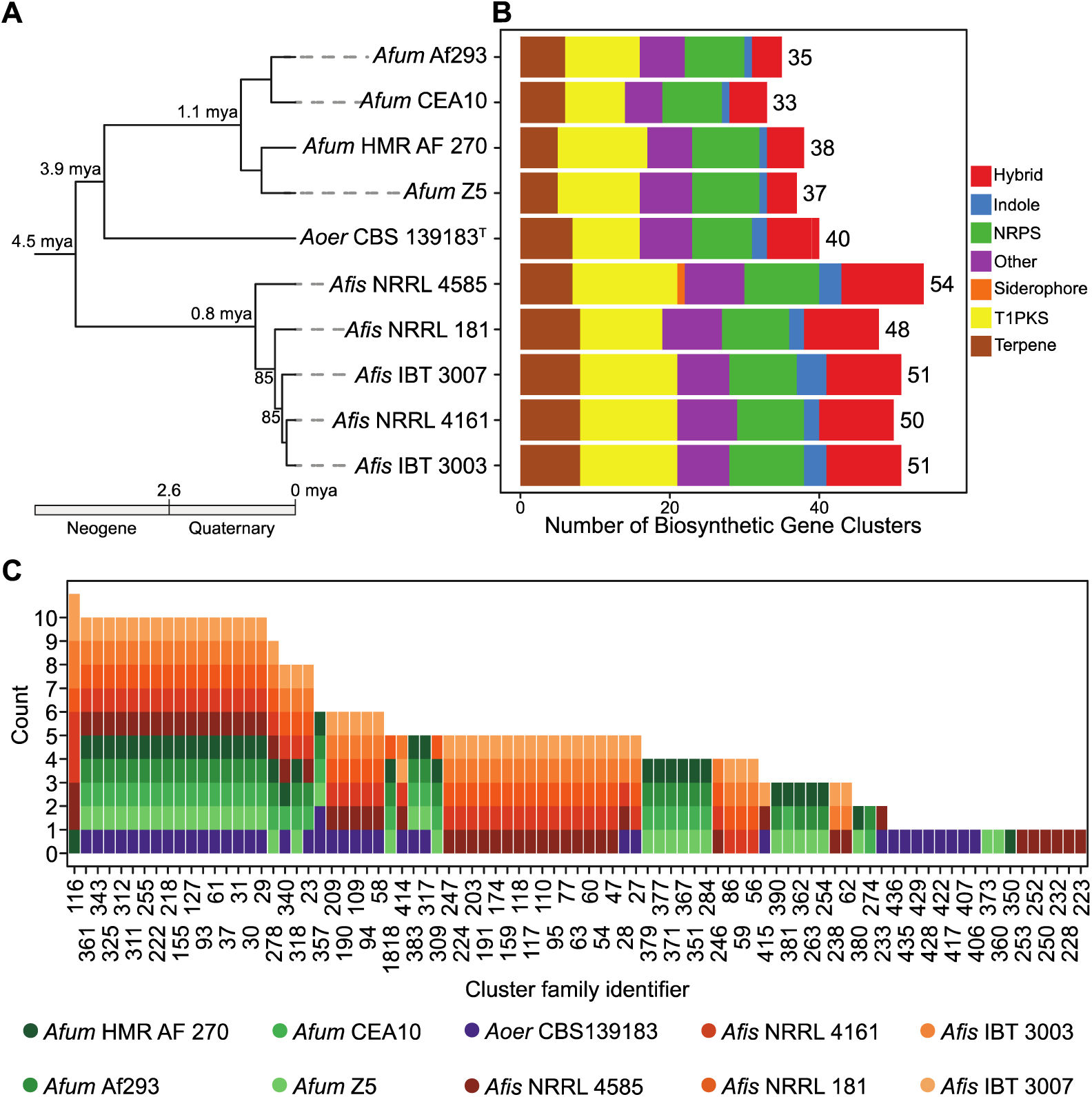
Diverse genetic repertoire of biosynthetic gene clusters and extensive presence and absence polymorphisms between and within species. (A) Genome-scale phylogenomic confirms *A. oerlinghausenensis* is the closest relative to *A. fumigatus*. Relaxed molecular clock analyses suggest *A. fumigatus, A. oerlinghausenensis*, and *A. fischeri* diverged from one another during the Neogene geologic period. Bipartition support is depicted for internodes that did not have full support. (B) *A. fumigatus* harbors the lowest number of BGCs compared to its two closest relatives. (C) Network-based clustering of BGCs into cluster families reveal extensive cluster presence and absence polymorphisms between species and strains. Cluster family identifiers are depicted on the x-axis; the number of strains represented in a cluster family are shown on the y-axis; the colors refer to a single strain from each species. Genus and species names are written using the following abbreviations: *Afum*: *A. fumigatus*; *Aoer*: *A. oerlinghausenensis*; *Afis*: *A. fischeri*.

Examination of the total number of predicted BGCs revealed that *A. fischeri* has the largest BGC repertoire. Among *A. fumigatus, A. oerlinghausenensis*, and *A. fischeri*, we predicted 50.80 ± 2.17, 40, 35.75 ± 2.22 BGCs, respectively, and found they spanned diverse biosynthetic classes (e.g., polyketides, non-ribosomal peptides, terpenes, etc.) (Fig. 1B). Network-based clustering of BGCs into cluster families (or groups of homologous BGCs) resulted in qualitatively similar networks when we used moderate similarity thresholds (or edge cut-off values; Fig. S3). Using a (moderate) similarity threshold of 0.5, we inferred 88 cluster families of putatively homologous BGCs (Fig. 1C).

Examination of BGCs revealed extensive presence and absence polymorphisms within and between species. We identified 17 BGCs that were present in all 10 *Aspergillus* genomes including the hexadehydroastechrome (HAS) BGC (cluster family 311 or CF311), the neosartoricin BGC (CF61), and other putative BGCs likely encoding unknown products (Fig. S4A; Table S1). In contrast, we identified 18 BGCs found in single strains, which likely encode unknown products. Between species, similar patterns of broadly present and species-specific BGCs were observed. For example, we identified 18 BGCs that were present in at least one strain across all species; in contrast, *A. fumigatus, A. oerlinghausenensis*, and *A. fischeri* had 16, eight, and 27 BGCs present in at least one strain but absent from the other species, respectively. These results suggest each species has a largely distinct repertoire of BGCs.

Examination of shared BGCs across species revealed *A. oerlinghausenensis* CBS139183^T^ and *A. fischeri* shared more BGCs with each other than either did with *A. fumigatus*. Surprisingly, we found ten homologous BGCs between *A. oerlinghausenensis* CBS 139183^T^ and *A. fischeri* but only three homologous BGCs shared between *A. fumigatus* and *A. oerlinghausenensis* CBS 139183^T^ (Fig. 2A; Fig. S4B) even though *A. oerlinghausenensis* is more closely related to *A. fumigatus* than to *A. fischeri* (Fig. 1A). BGCs shared by *A. oerlinghausenensis* CBS 139183^T^ and *A. fischeri* were uncharacterized while BGCs present in both *A. fumigatus* and *A. oerlinghausenensis* CBS 139183^T^ included those that encode fumigaclavine and fumagillin/pseurotin. Lastly, to associate each BGC with a secondary metabolite in *A. fumigatus* Af293, we cross referenced our list with a publicly available one (Table S2). Importantly, all known *A. fumigatus* Af293 BGCs were represented in our analyses.

**Figure 2.**
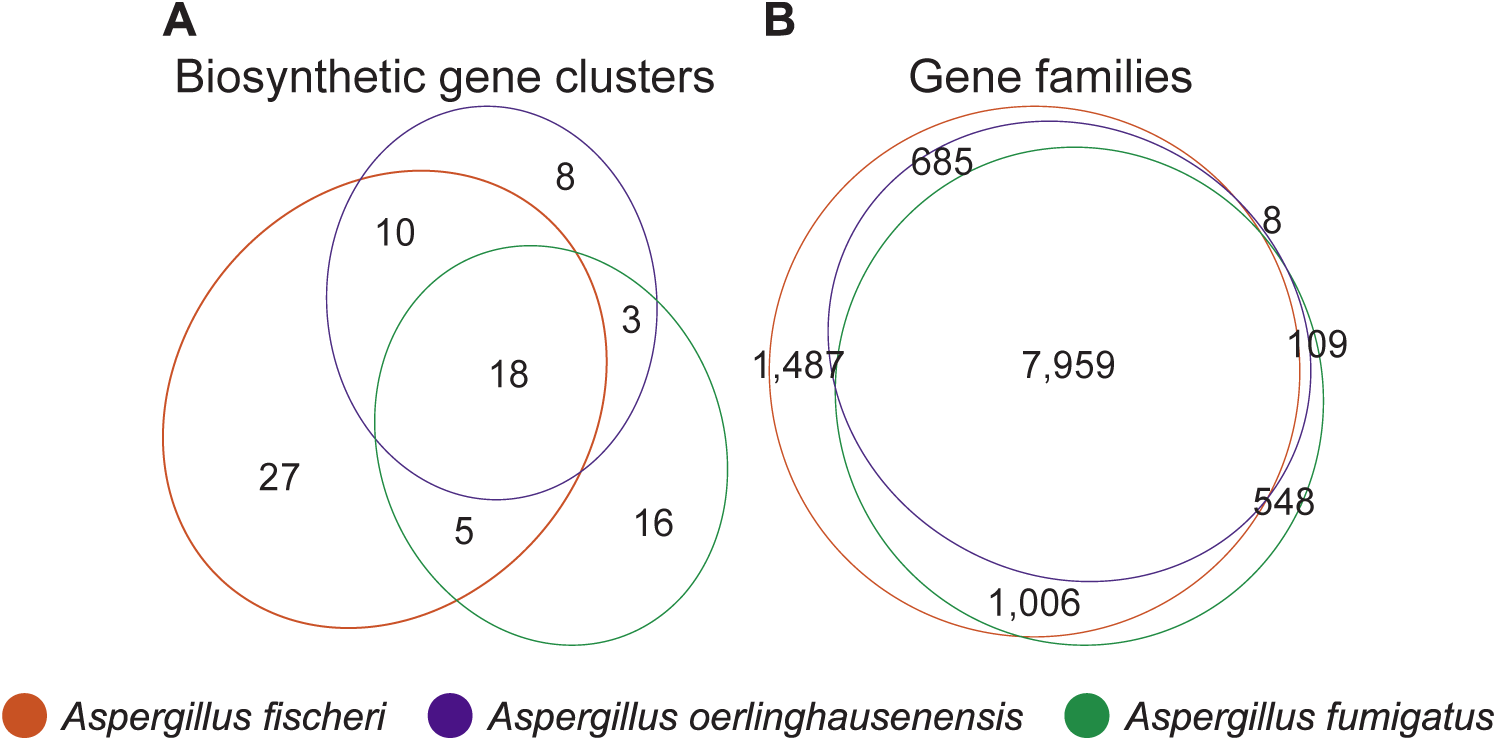
*Aspergillus oerlinghausenensis* shares more gene families and BGCs with *A. fischeri* than *A. fumigatus*. (A) Euler diagram showing species-level shared BGCs. (B) Euler diagram showing species-level shared gene families. In both diagrams, *A. oerlinghausenensis* shares more gene families or BGCs with *A. fischeri* than *A. fumigatus* despite a closer evolutionary relationship.

More broadly, examination of shared and species-specific gene families revealed that *A. oerlinghausenensis* does not have many species-specific gene families and shares more gene families with *A. fischeri* than *A. fumigatus* (Fig. 2B). Specifically, we noted that *A. oerlinghausenensis* CBS 139183^T^ has only eight species-specific gene families; in contrast, *A. fischeri* and *A. fumigatus* have 1,487 and 548 species-specific gene families, respectively.

Despite a closer evolutionary relationship between *A. oerlinghausenensis* and *A. fumigatus*, we found *A. oerlinghausenensis* shares more gene families with *A. fischeri* compared to *A. fumigatus* (685 and 109, respectively) suggestive of extensive gene loss in the *A. fumigatus* stem lineage. Lastly, we observed strain heterogeneity in gene family presence and absence within both *A. fumigatus* and *A. fischeri* (Fig. S5).

### Within and between species variation in secondary metabolite profiles of *A. fumigatus* and its closest relatives

To gain insight into variation in secondary metabolite profiles within and between species, we profiled *A. fumigatus* strains Af293, CEA10, and CEA17 (a *pyrG1/URA3* derivative of CEA10), *A. fischeri* strains NRRL 181, NRRL 4585, and NRRL 4161, and *A. oerlinghausenensis* CBS 139183^T^ for secondary metabolites. Specifically, we used three different procedures, including the isolation and structure elucidation of metabolites, where possible, followed by two different metabolite profiling procedures that use mass spectrometry techniques. Altogether, we isolated and characterized 19 secondary metabolites; seven from *A. fumigatus*, two from *A. oerlinghausenensis*, and ten from *A. fischeri* (Fig. S6). These products encompassed a wide diversity of secondary metabolite classes, such as those derived from polyketide synthases, non-ribosomal peptide-synthetases, terpene synthases and mixed biosynthesis enzymes.

To characterize the secondary metabolites biosynthesized that were not produced in high enough quantity for structural identification through traditional isolation methods, we employed “dereplication” mass spectrometry protocols specific to natural products research on all tested strains at both 30°C and 37°C (see supporting information, dereplication example; figshare: 10.6084/m9.figshare.12055503) (El-Elimat et al., 2013; Ito and Masubuchi, 2014; Gaudêncio and Pereira, 2015; Hubert et al., 2017). We found an overlap of secondary metabolites between strains of the same species (Table S3); for example, monomethylsulochrin (**3**) was isolated from *A. fumigatus* Af293, but through metabolite profiling, its spectral features were noted also in *A. fumigatus* strains CEA10 and CEA17. We identified metabolites that were biosynthesized by only one species; for example, pseurotin A (**4**) was solely present in *A. fumigatus* strains. Furthermore, we also found an overlap of several secondary metabolites across species, such as fumagillin (**6**), which was biosynthesized by *A. fumigatus* and *A. oerlinghausenensis*, and fumitremorgin B (**16**), which was biosynthesized by strains of both *A. oerlinghausenensis* and *A. fischeri.* Together, these analyses suggest that closely related *Aspergillus* species and strains exhibit variation both within as well as between species in the secondary metabolites produced. To further facilitate comparisons of secondary metabolite profiles within and between species, we used the 1,920 features (i.e., unique *m/z* – retention time pairs) that were identified from all strains at all temperatures (Fig. 3A), to perform Principal Components Analysis (PCA) (Fig. S7) and hierarchical clustering (Fig. 3B). The PCA plots and hierarchical clustering of the chromatograms at both 37°C and 30°C (Fig. 3B, S7B, and S7C) indicated that the three *A. fischeri* strains and *A. oerlinghausenensis* were the most chemically similar to each other. When we generated PCA plots from only the compounds that could be isolated from each culture (44 features that represent the 19 isolated compounds) at both 37°C (Fig. S7D) and 30°C (Fig. S7E), the chemical similarities between *A. oerlinghausenensis* and *A. fischeri* were even more evident, with this clustering showing similar results to the total feature dendrogram (Fig. 3B). These data suggest that there are more similarities among the secondary metabolites that are biosynthesized in higher abundance. While the clustering of *A. oerlinghausenensis* CBS 139183^T^ and *A. fischeri* NRRL 181 is conserved, these strains were also shown to be similar to *A. fumigatus* CEA10 (Fig. S7D-E). These combined results suggest that at lower temperatures, such as 30°C, there is a more varied response in how BGCs are being utilized, leading to a more diverse production of chemical compounds.

**Figure 3.**
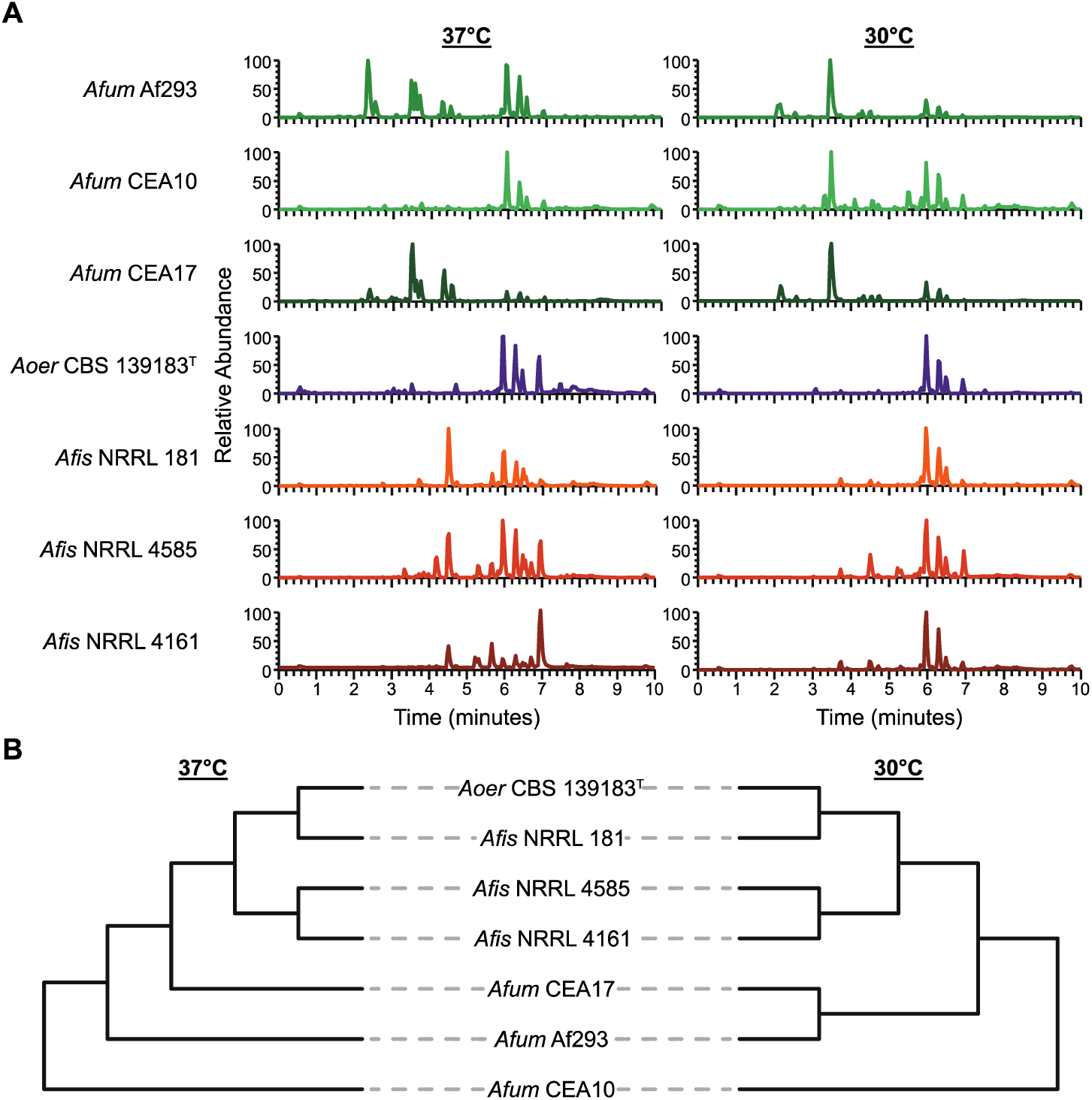
*A. oerlinghausenensis* and *A. fischeri* have more similar secondary metabolite profiles than *A. fumigatus*. (A) UPLC-MS chromatograms of secondary metabolite profiles of *A. fumigatus* and its closest relatives, *A. oerlinghausenensis* and *A. fischeri* at 37°C and 30°C (left and right, respectively). (B) Hierarchical clustering of chromatograms reveals *A. oerlinghausenensis* clusters with *A. fischeri* and not its closest relative, *A. fumigatus* at 37°C and 30°C (left and right, respectively).

In summary, our chemical analyses suggest that the secondary metabolite profiles of *A. oerlinghausenensis* and *A. fischeri* are more similar to each other than to *A. fumigatus* (Fig. 3B and S7B-E). This finding is surprising because phylogenetic analysis indicates that *A. oerlinghausenensis* is more closely related to *A. fumigatus* than it is to *A. fischeri* (Fig. 1A). However, the similarity of secondary metabolite profiles of *A. oerlinghausenensis* and *A. fischeri* is consistent with our finding that the genome of *A. oerlinghausenensis* shares higher numbers of BGCs and gene families with *A. fischeri* than with *A. fumigatus* (Fig. 2). Similarly, clustering patterns in secondary metabolite-based plots (Fig. S7B-E) resemble those of BGC-based plots (Fig. S7A), suggesting that the observed similarities in the chemotypes of *A. oerlinghausenensis* and *A. fischeri* are broadly reflected in their metabolism-associated genotypes.

### Conservation and divergence among biosynthetic gene clusters implicated in *A. fumigatus* pathogenicity

Secondary metabolites are known to play a role in *A. fumigatus* virulence (Raffa and Keller, 2019). We therefore conducted a focused examination of specific *A. fumigatus* BGCs and secondary metabolites that have been previously implicated in the organism’s ability to cause human disease (Table 1). We found varying degrees of conservation and divergence that were associated with the absence or presence of a secondary metabolite. Among conserved BGCs that were also associated with conserved secondary metabolite production, we highlight the mycotoxins gliotoxin and fumitremorgin. Interestingly, we note that only *A. fischeri* strains synthesized verruculogen, a secondary metabolite that is implicated in human disease and is encoded by the fumitremorgin BGC (Khoufache et al., 2007; Kautsar et al., 2019). Among divergent BGCs that were associated with the absence of a secondary metabolite, we highlight the trypacidin and fumagillin/pseurotin secondary metabolites. We found that nonpathogenic close relatives of *A. fumigatus* produced some but not all mycotoxins, which provides novel insight into the unique cocktail of secondary metabolites biosynthesized by *A. fumigatus*.

#### Gliotoxin

Gliotoxin is a highly toxic compound and known virulence factor in *A. fumigatus* (Sugui et al., 2007). Nearly identical BGCs encoding gliotoxin are present in all pathogenic (*A. fumigatus*) and nonpathogenic (*A. oerlinghausenensis* and *A. fischeri*) strains examined (Fig. 4). Additionally, we found that all examined strains synthesized bisdethiobis(methylthio)gliotoxin a derivative from dithiogliotoxin, involved in the down-regulation of gliotoxin biosynthesis (Dolan et al., 2014), one of the main mechanisms of gliotoxin resistance in *A. fumigatus* (Kautsar et al., 2019).

**Figure 4.**
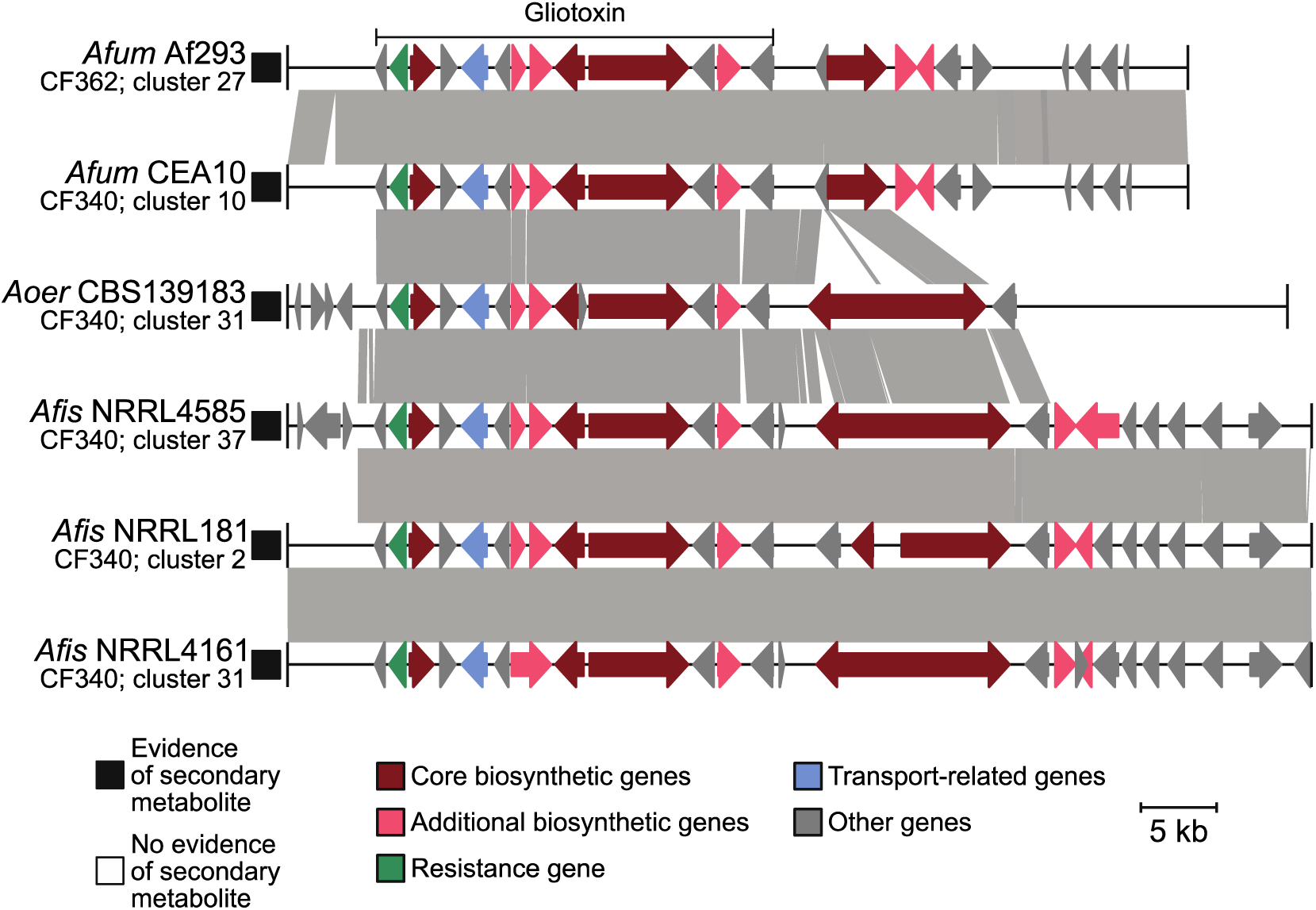
Conservation in the gliotoxin BGC correlates with conserved production of gliotoxin analogs in *A. fumigatus* and nonpathogenic close relatives. Microsynteny analysis reveals a high degree of conservation in the BGC encoding gliotoxin across all isolates. The known gliotoxin gene cluster boundary is indicated above the *A. fumigatus* Af293 BGC. Black and white squares correspond to the presence or absence of the associated secondary metabolite, respectively. Genes are drawn as arrows with orientation indicated by the direction of the arrow. Gene function is indicated by gene color. Genus and species names are written using the following abbreviations: *Afum*: *A. fumigatus*; *Aoer*: *A. oerlinghausenensis*; *Afis*: *A. fischeri*. Below each genus and species abbreviation is the cluster family each BGC belongs to and their cluster number.

#### Fumitremorgin and Verruculogen

Similarly, there is a high degree of conservation in the BGC that encodes fumitremorgin across all strains (Fig. 5). Fumitremorgins have known antifungal activity, are lethal to brine shrimp, and are implicated in inhibiting mammalian proteins responsible for resistance to anticancer drugs in mammalian cells (Raffa and Keller, 2019). We found that conservation in the fumitremorgin BGC is associated with the production of fumitremorgins in all isolates examined. The fumitremorgin BGC is also responsible for the production of verruculogen, which is implicated to aid in *A. fumigatus* pathogenicity by changing the electrophysical properties of human nasal epithelial cells (Khoufache et al., 2007). Interestingly, we found that only *A. fischeri* strains produced verruculogen under the conditions we analyzed.

**Figure 5.**
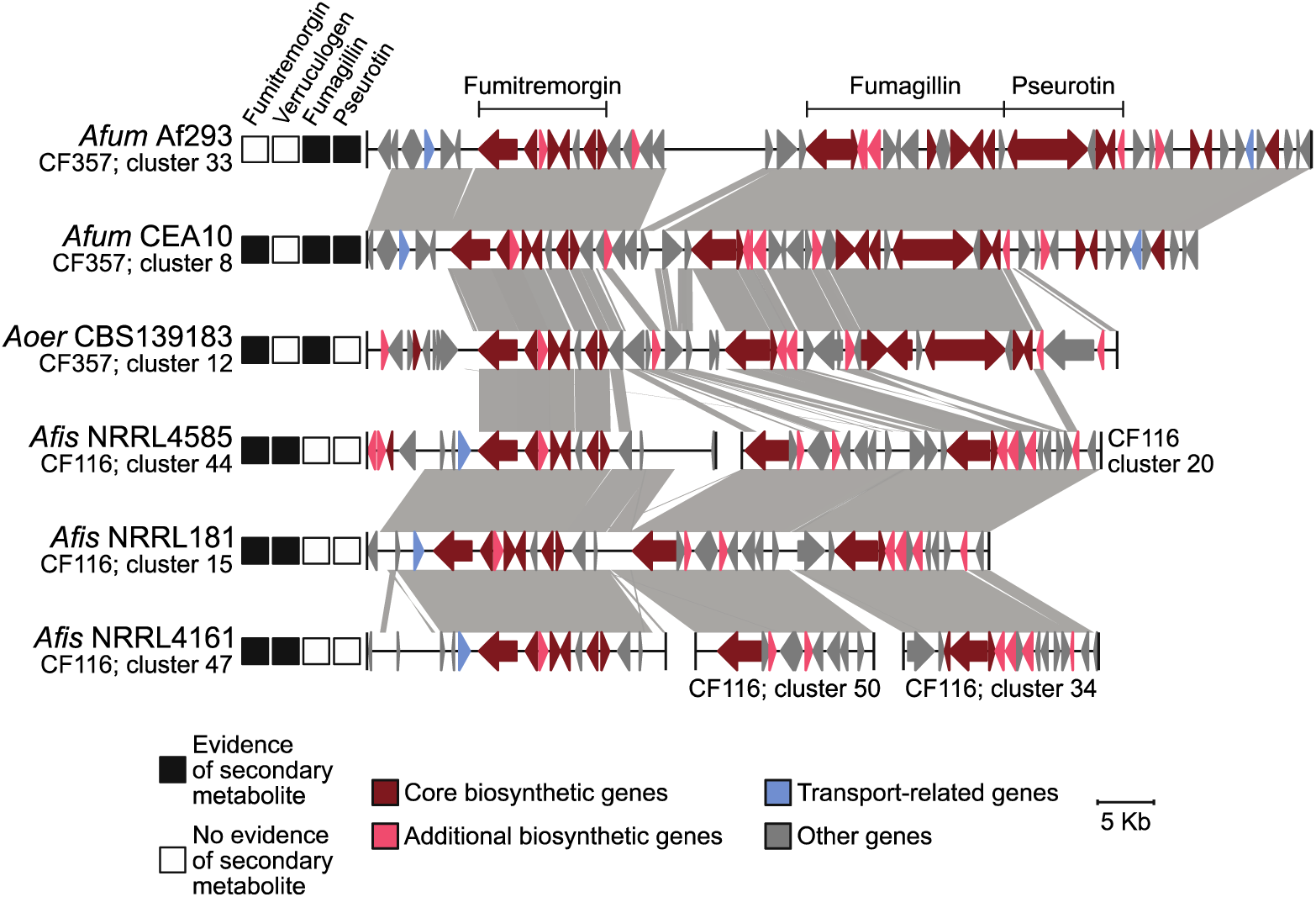
Conservation and divergence in the locus encoding the fumitremorgin and intertwined fumagillin/pseurotin BGCs. Microsynteny analysis reveals conservation in the fumitremorgin BGC across all isolates. Interestingly, only *A. fischeri* strains synthesize verruculogen, a secondary metabolite also biosynthesized by the fumitremorgin BGC. In contrast, the intertwined fumagillin/pseurotin BGCs are conserved between *A. fumigatus* and *A. oerlinghausenensis* but divergent in *A. fischeri*. BGC conservation and divergence is associated with the presence and absence of a secondary metabolite, respectively. The same convention used in Fig. 4 is used to depict evidence of a secondary metabolite, represent genes and broad gene function, genus and species abbreviations, and BGC cluster families and cluster numbers.

#### Trypacidin

Examination of the trypacidin BGC, which encodes a spore-borne and cytotoxic secondary metabolite, revealed a conserved cluster found in four pathogenic and nonpathogenic strains: *A. fumigatus* Af293, *A. fumigatus* CEA10, *A. oerlinghausenensis* CBS 139183^T^, and *A. fischeri* NRRL 181 (Fig. S8). Furthermore, we found that three of these four isolates (except *A. fischeri* NRRL 181) biosynthesized a trypacidin analog, monomethylsulochrin. Examination of the microsynteny of the trypacidin BGC revealed that it was conserved across all four genomes with the exception *A. fischeri* NRRL l81, which lacked a RING (Really Interesting New Gene) finger gene. Interestingly, RING finger proteins can mediate gene transcription (Poukka et al., 2000). We confirmed the absence of the RING finger protein by performing a sequence similarity search with the *A. fumigatus* Af293 RING finger protein (AFUA_4G14620; EAL89333.1) against the *A. fischeri* NRRL 181 genome. In the homologous locus in *A. fischeri*, we found no significant blast hit for the first 23 nucleotides of the RING finger gene suggestive of pseudogenization. Taken together, we hypothesize that presence/absence polymorphisms or a small degree of sequence divergence between otherwise homologous BGCs may be responsible for the presence or absence of a toxic secondary metabolite in *A. fischeri*.

#### Fumagillin/pseurotin

Examination of the intertwined fumagillin/pseurotin BGCs revealed that fumagillin has undergone substantial sequence divergence and that pseurotin is absent from strains of *A. fischeri*. The fumagillin/pseurotin BGCs are under the same regulatory control (Wiemann et al., 2013) and biosynthesize secondary metabolites that cause cellular damage during host infection (fumagillin (Guruceaga et al., 2019)) and inhibit immunoglobulin E production (pseurotin (Ishikawa et al., 2009)). Microsynteny of the fumagillin BGC reveals high sequence conservation between *A. fumigatus* and *A. oerlinghausenensis*; however, sequence divergence was observed between *A. oerlinghausenensis* and *A. fischeri* (Fig. 5). Accordingly, fumagillin production was only observed in *A. fumigatus* and *A. oerlinghausenensis* and not in *A. fischeri*. Similarly, the pseurotin BGC is conserved between *A. fumigatus* and *A. oerlinghausenensis*. Rather than sequence divergence, no sequence similarity was observed in the region of the pseurotin cluster in *A. fischeri*, which may be due to an indel event. Accordingly, no pseurotin production was observed among *A. fischeri* strains. Despite sequence conservation between *A. fumigatus* and *A. oerlinghausenensis*, no evidence of pseurotin biosynthesis was observed in *A. oerlinghausenensis*, which suggests regulatory decoupling of the intertwined fumagillin/pseurotin BGC. Altogether, these results show a striking correlation between sequence divergence and the production (or absence) of secondary metabolites implicated in human disease among *A. fumigatus* and nonpathogenic closest relatives.

## Discussion

*Aspergillus fumigatus* is a major fungal pathogen nested within a clade (known as section *Fumigati*) of at least 60 other species, the vast majority of which are nonpathogenic (Steenwyk et al., 2019; Rokas et al., 2020a). Currently, it is thought that the ability to cause human disease evolved multiple times among species in section *Fumigati* (Rokas et al., 2020a). Secondary metabolites contribute to the success of the major human pathogen *A. fumigatus* in the host environment (Raffa and Keller, 2019) and are therefore “cards” of virulence (Casadevall, 2007; Knowles et al., 2020). However, whether the closest relatives of *A. fumigatus, A. oerlinghausenensis* and *A. fischeri*, both of which are nonpathogenic, biosynthesize secondary metabolites implicated in the ability of *A. fumigatus* to cause human disease remained largely unknown. By examining genomic and chemical variation between and within *A. fumigatus* and its closest nonpathogenic relatives, we identified both conservation and divergence (including within species heterogeneity) in BGCs and secondary metabolite profiles (Fig. 1-5, S4, S6-9; Table 1, S1, S3). Examples of conserved BGCs and secondary metabolites include the major virulence factor, gliotoxin (Fig. 4), as well as several others (Fig. 5, S8; Table 1, S1, S3); examples of BGC and secondary metabolite heterogeneity or divergence include pseurotin, fumagillin, and several others (Fig. 5; Table 1, S1, S3). Lastly, we found that the fumitremorgin BGC, which biosynthesizes fumitremorgin in all three species, is also associated with verruculogen biosynthesis in *A. fischeri* strains (Fig. 5).

One of the surprising findings of our study was that although *A. oerlinghausenensis* and *A. fumigatus* are evolutionarily more closely related to each other than to *A. fischeri* (Fig. 1), *A. oerlinghausenensis* and *A. fischeri* appear to be more similar to each other than to *A. fumigatus* in BGC composition, gene family content, and secondary metabolite profiles. The power of pathogen-nonpathogen comparative genomics is best utilized when examining closely related species (Fedorova et al., 2008; Jackson et al., 2011; Moran et al., 2011; Mead et al., 2019a; Rokas et al., 2020a). By sequencing genomes from the closest known nonpathogenic relatives of *A. fumigatus*, including the genome of the closest species relative *A. oerlinghausenensis* and additional strains of *A. fischeri*, we provide a powerful resource to study the evolution of *A. fumigatus* pathogenicity.

Our finding that *A. oerlinghausenensis* and *A. fischeri* shares more gene families and BGCs with each other than they do with *A. fumigatus* (Fig. 1C, 2, S4, S5, S9) suggests that the evolutionary trajectory of the *A. fumigatus* ancestor was marked by gene loss. We hypothesize that there were two rounds of gene family and BGC loss in the *A. fumigatus* stem lineage: (1) gene families and BGCs were lost in the common ancestor of *A. fumigatus* and *A. oerlinghausenensis* and (2) additional losses occurred in the *A. fumigatus* ancestor. In addition to losses, we note that 548 and 16 gene families and BGCs are unique to *A. fumigatus*, which may have resulted from genetic innovation (e.g., *de novo* gene formation) or unique gene family and BGC retention (Fig. 2, S9). In line with the larger number of shared BGCs between *A. oerlinghausenensis* and *A. fischeri*, we found their secondary metabolite profiles were also more similar (Fig. 3, S7). Notably, the evolutionary rate of the internal branch leading to the *A. fumigatus* common ancestor is much higher than those in the rest of the branches in our genome-scale phylogeny (Fig. S2B), suggesting that the observed gene loss and gene gain / retention events specific to *A. fumigatus* may be part of a wider set of evolutionary changes in the *A. fumigatus* genome. More broadly, these results suggest that comparisons of the pathogen *A. fumigatus* against either the non-pathogen *A. oerlinghausenensis* (this manuscript) or the non-pathogen *A. fischeri* ((Mead et al., 2019a; Knowles et al., 2020) and this manuscript) will both be instructive in understanding the evolution of *A. fumigatus* pathogenicity.

When studying *Aspergillus* pathogenicity, it is important to consider any genetic and phenotypic heterogeneity between strains of a single species (Kowalski et al., 2016, 2019; Keller, 2017; Ries et al., 2019; Bastos et al., 2020; Santos et al., 2020). Our finding of strain heterogeneity among gene families, BGCs, and secondary metabolites in *A. fumigatus* and *A. fischeri* (Fig. 1-3, S4, S5, S7, S9) suggests considerable strain-level diversity in each species. For example, we found secondary metabolite profile strain heterogeneity was greater in *A. fumigatus* than *A. fischeri* (Fig. S7B-E). These results suggest that strain specific secondary metabolite profiles may play a role in variation of pathogenicity among *A. fumigatus* strains. More broadly, our finding supports the hypothesis that strain-level diversity is an important parameter when studying pathogenicity (Kowalski et al., 2016, 2019; Keller, 2017; Ries et al., 2019; Bastos et al., 2020; Santos et al., 2020).

Secondary metabolites contribute to *A. fumigatus* virulence through diverse processes including suppressing the human immune system and damaging tissues (Table 1). Interestingly, we found that the nonpathogens *A. oerlinghausenensis* and *A. fischeri* produced several secondary metabolites implicated in the ability of *A. fumigatus* human disease, such gliotoxin, trypacidin, verruculogen, and others (Fig. 4, 5, S8; Table 1, S3). Importantly, our work positively identified secondary metabolites for many structural classes implicated in a previous taxonomic study (Samson et al., 2007). These results suggest that several of the secondary metabolism-associated cards of virulence present in *A. fumigatus* are conserved in closely related nonpathogens (summarized in Fig. 6). Interestingly, disrupting the ability of *A. fumigatus* to biosynthesize gliotoxin attenuates but does not abolish virulence (Sugui et al., 2007; Dagenais and Keller, 2009; Keller, 2017), whereas disruption of the ability of *A. fischeri* NRRL 181 to biosynthesize secondary metabolites, including gliotoxin, does not appear to influence virulence (Knowles et al., 2020). Our findings, together with previous studies, support the hypothesis that individual secondary metabolites are “cards” of virulence in a larger “hand” that *A. fumigatus* possesses.

**Figure 6.**
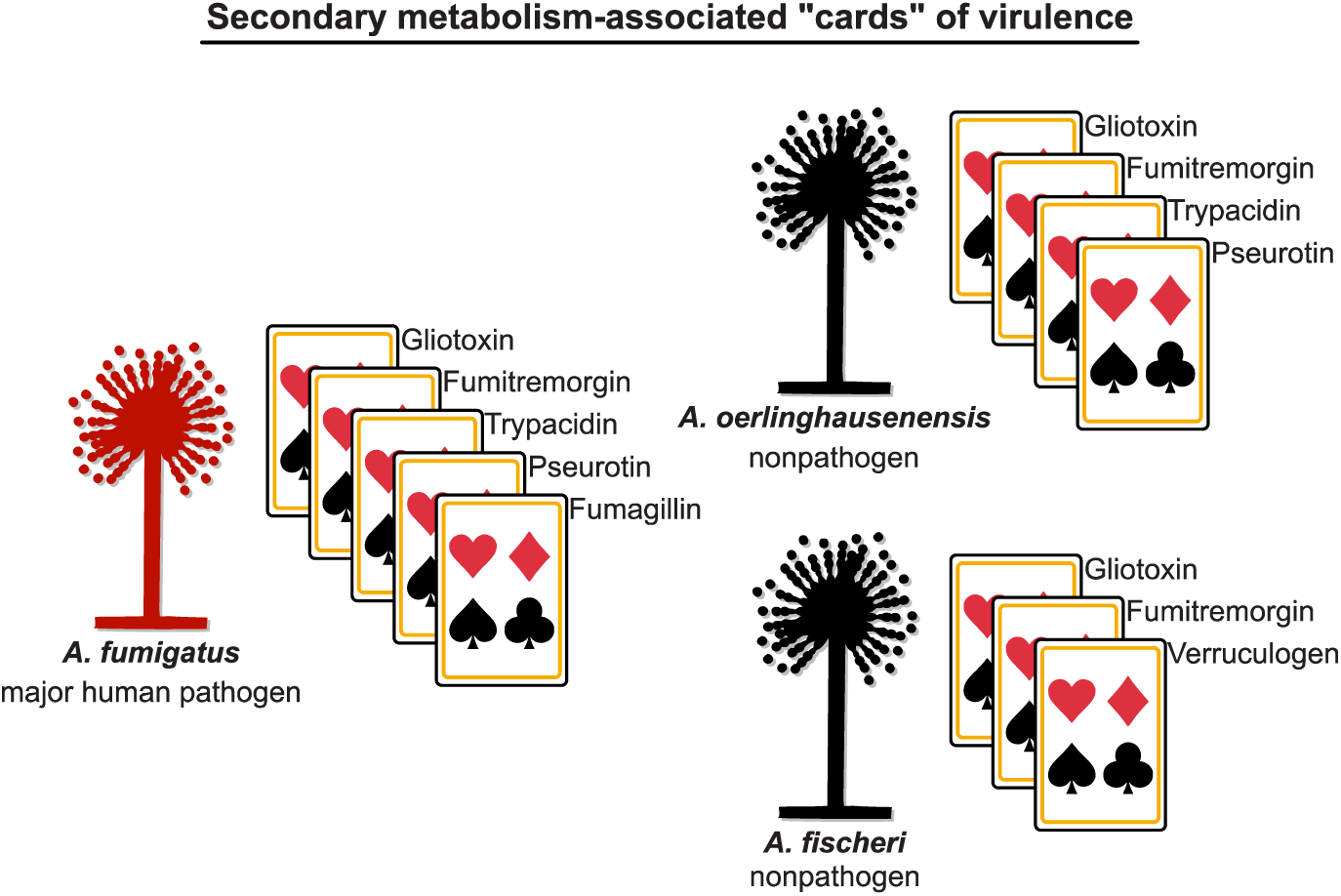
Secondary metabolism-associated “cards” of virulence among *A. fumigatus* and close relatives. Secondary metabolites contribute to the “hand of cards”’ that enable *A. fumigatus* to cause disease. Here, we show that the nonpathogenic closest relatives of *A. fumigatus* possess a subset of the *A. fumigatus* secondary metabolism-associated cards of virulence. We hypothesize that the unique combination of cards of *A. fumigatus* contributes to its pathogenicity and that the cards in *A. oerlinghausenensis* and *A. fischeri* (perhaps in combination with other non-secondary-metabolism-associated cards, such as thermotolerance) are insufficient to cause disease. Pathogenic and nonpathogenic species are shown in red and black, respectively. Cartoons of *Aspergillus* species were obtained from WikiMedia Commons (source: M. Piepenbring) and modified in accordance with the Creative Commons Attribution-Share Alike 3.0 Unported license (https://creativecommons.org/licenses/by-sa/3.0/deed.en).

## Methods

### Strain acquisition, DNA extraction, and sequencing

Two strains of *Aspergillus fischeri* (NRRL 4161 and NRRL 4585) were acquired from the Northern Regional Research Laboratory (NRRL) at the National Center for Agricultural Utilization Research in Peoria, Illinois, while one strain of *Aspergillus oerlinghausenensis* (CBS 139183^T^) was acquired from Westerdijk Fungal Biodiversity Institute, Utrecht, The Netherlands. These strains were grown in 50 ml of liquid yeast extract soy peptone dextrose (YESD) medium. After approximately seven days of growth on an orbital shaker (100 rpm) at room temperature, the mycelium was harvested by filtering the liquid media through a Corning®, 150 ml bottle top, 0.22µm sterile filter and washed with autoclaved distilled water. All subsequent steps of DNA extraction from the mycelium were performed following protocols outlined previously (Mead et al., 2019b). The genomic DNA from these three strains was sequenced using a NovaSeq S4 at the Vanderbilt Technologies for Advanced Genomes facility (Nashville, Tennessee, US) using paired-end sequencing (150 bp) strategy with the Illumina TruSeq library kit.

### Genome assembly, quality assessment, and annotation

To assemble and annotate the three newly sequenced genomes, we first quality-trimmed raw sequence reads using Trimmomatic, v0.36 (Bolger et al., 2014) using parameters described elsewhere (ILLUMINACLIP:TruSeq3-PE.fa:2:30:10, leading:10, trailing:10, slidingwindow:4:20, minlen:50) (Steenwyk and Rokas, 2017). The resulting paired and unpaired quality-trimmed reads were used as input to the SPAdes, v3.11.1 (Bankevich et al., 2012), genome assembly algorithm with the ‘careful’ parameter and the ‘cov-cutoff’ set to ‘auto’. We evaluated the quality of our newly assembled genomes, using metrics based on continuity of assembly and gene-content completeness. To evaluate genome assemblies by scaffold size, we calculated the N50 of each assembly (or the shortest contig among the longest contigs that account for 50% of the genome assembly’s length) (Yandell and Ence, 2012). To determine gene-content completeness, we implemented the BUSCO, v2.0.1 (Waterhouse et al., 2018), pipeline using the ‘genome’ mode. In this mode, the BUSCO pipeline examines assembly contigs for the presence of near-universally single copy orthologous genes (hereafter referred to as BUSCO genes) using a predetermined database of orthologous genes from the OrthoDB, v9 (Waterhouse et al., 2013). We used the OrthoDB database for Pezizomycotina (3,156 BUSCO genes). Each BUSCO gene is determined to be present in a single copy, as duplicate sequences, fragmented, or missing. Our analyses indicate the newly sequenced and assembled genomes have high gene-content completeness and assembly continuity (average percent presence of BUSCO genes: 98.80 ± 0.10%; average N50: 451,294.67 ± 9,696.11; Fig. S1). These metrics suggest these genomes are suitable for comparative genomic analyses.

To predict gene boundaries in the three newly sequenced genomes, we used the MAKER, v2.31.10, pipeline (Holt and Yandell, 2011) which, creates consensus predictions from the collective evidence of multiple *ab initio* gene prediction software. Specifically, we created consensus predictions from SNAP, v2006-07-28 (Korf, 2004), and AUGUSTUS, v3.3.2 (Stanke and Waack, 2003), after training each algorithm individually on each genome. To do so, we first ran MAKER using protein evidence clues from five different publicly available annotations of *Aspergillus* fungi from section *Fumigati*. Specifically, we used protein homology clues from *A. fischeri* NRRL 181 (GenBank accession: GCA_000149645.2), *A. fumigatus* Af293 (GenBank accession: GCA_000002655.1), *Aspergillus lentulus* IFM 54703 (GenBank accession: GCA_001445615.1), *Aspergillus novofumigatus* IBT 16806 (GenBank accession: GCA_002847465.1), and *Aspergillus udagawae* IFM 46973 (GenBank accession: GCA_001078395.1). The resulting gene predictions were used to train SNAP. MAKER was then rerun using the resulting training results. Using the SNAP trained gene predictions, we trained AUGUSTUS. A final set of gene boundary predictions were obtained by rerunning MAKER with the training results from both SNAP and AUGUSTUS.

To supplement our data set of newly sequenced genomes, we obtained publicly available ones. Specifically, we obtained genomes and annotations for *A. fumigatus* Af293 (GenBank accession: GCA_000002655.1), *A. fumigatus* CEA10 (strain synonym: CBS 144.89 / FGSC A1163; GenBank accession: GCA_000150145.1), *A. fumigatus* HMR AF 270 GenBank accession: GCA_002234955.1), *A. fumigatus* Z5 (GenBank accession: GCA_001029325.1), *A. fischeri* NRRL 181 (GenBank accession: GCA_000149645.2). We also obtained assemblies of the recently published *A. fischeri* genomes for strains IBT 3003 and IBT 3007 (Zhao et al., 2019) which, lacked annotations. We annotated the genome of each strain individually using MAKER with the SNAP and AUGUSTUS training results from a close relative of both strains, *A. fischeri* NRRL 4161. Altogether, our final data set contained a total of ten genome from three species: four *A. fumigatus* strains, one *A. oerlinghausenensis* strain, and five *A. fischeri* strains.

### Maximum likelihood phylogenetics and Bayesian estimation of divergence times

To reconstruct the evolutionary history among the ten *Aspergillus* genomes, we implemented a recently developed pipeline (Steenwyk et al., 2019) which, relies on the concatenation-approach to phylogenomics (Rokas et al., 2003) and has been successfully used in reconstructing species-level relationships among *Aspergillus* and *Penicillium* fungi (Bodinaku et al., 2019; Steenwyk et al., 2019). The first step in the pipeline is to identify single copy orthologous genes in the genomes of interest which, are ultimately concatenated into a larger phylogenomic data matrix. To identify single copy BUSCO genes across all ten *Aspergillus* genomes, we used the BUSCO pipeline with the Pezizomycotina database as described above. We identified 3,041 BUSCO genes present at a single copy in all ten *Aspergillus* genomes and created multi-FASTA files for each BUSCO gene that contained the protein sequences for all ten taxa. The protein sequences of each BUSCO gene were individually aligned using Mafft, v7.4.02 (Katoh and Standley, 2013), with the same parameters as described elsewhere (Steenwyk et al., 2019). Nucleotide sequences were then forced onto the protein sequence alignments using a custom Python, v3.5.2 (https://www.python.org/), script with BioPython, v1.7 (Cock et al., 2009). The resulting codon-based alignments were trimmed using trimAl, v1.2.rev59 (Capella-Gutierrez et al., 2009), with the ‘gappyout’ parameter. The resulting trimmed nucleotide alignments were concatenated into a single matrix of 5,602,272 sites and was used as input into IQ-TREE, v1.6.11 (Nguyen et al., 2015). The best-fitting model of substitutions for the entire matrix was determined using Bayesian information criterion values (Kalyaanamoorthy et al., 2017). The best-fitting model was a general time-reversible model with empirical base frequencies that allowed for a proportion of invariable sites and a discrete Gamma model with four rate categories (GTR+I+F+G4) (Tavaré, 1986; Yang, 1994, 1996; Vinet and Zhedanov, 2011). To evaluate bipartition support, we used 5,000 ultrafast bootstrap approximations (Hoang et al., 2018).

To estimate divergence times among the ten *Aspergillus* genomes, we used the concatenated data matrix and the resulting maximum likelihood phylogeny from the previous steps as input to Bayesian approach implemented in MCMCTree from the PAML package, v4.9d (Yang, 2007). First, we estimated the substitution rate across the data matrix using a “GTR+G” model of substitutions (model = 7), a strict clock model, and the maximum likelihood phylogeny rooted on the clade of *A. fischeri* strains. We imposed a root age of 3.69 million years ago according to results from recent divergence time estimates of the split between *A. fischeri* and *A. fumigatus* (Steenwyk et al., 2019). We estimated the substitution rate to be 0.005 substitutions per one million years. Next, the likelihood of the alignment was approximated using a gradient and Hessian matrix. To do so, we used previously established time constraints for the split between *A. fischeri* and *A. fumigatus* (1.85 to 6.74 million years ago) (Steenwyk et al., 2019). Lastly, we used the resulting gradient and Hessian matrix, the rooted maximum likelihood phylogeny, and the concatenated data matrix to estimate divergence times using a relaxed molecular clock (model = 2). We specified the substitution rate prior based on the estimated substitution rate (rgene_gamma = 1 186.63). The ‘sigma2_gamma’ and ‘finetune’ parameters were set to ‘1 4.5’ and ‘1’, respectively. To collect a high-quality posterior probability distribution, we ran a total of 5.1 million iterations during MCMC analysis which, is 510 times greater than the minimum recommendations (Raftery and Lewis, 1995). Our sampling strategy across the 5.1 million iterations was to discard the first 100,000 results followed by collecting a sample every 500^th^ iteration until a total of 10,000 samples were collected.

### Identification of gene families and analyses of putative biosynthetic gene clusters

To identify gene families across the ten *Aspergillus* genomes, we used a Markov clustering approach. Specifically, we used OrthoFinder, v2.3.8 (Emms and Kelly, 2019). OrthoFinder first conducts a blast all-vs-all using the protein sequences of all ten *Aspergillus* genomes and NCBI’s Blast+, v2.3.0 (Camacho et al., 2009), software. After normalizing blast bit scores, genes are clustered into discrete orthogroups using a Markov clustering approach (van Dongen, 2000). We clustered genes using an inflation parameter of 1.5. The resulting orthogroups were used proxies for gene families.

To identify putative biosynthetic gene clusters (BGCs), we used the gene boundaries predictions from the MAKER software as input into antiSMASH, v4.1.0 (Weber et al., 2015). To identify homologous BGCs across the ten *Aspergillus* genomes, we used the software BiG-SCAPE, v20181005 (Navarro-Muñoz et al., 2020). Based on the Jaccard Index of domain types, sequence similarity among domains, and domain adjacency, BiG-SCAPE calculates a similarity metric between pairwise combinations of clusters where smaller values indicate greater BGC similarity. BiG-SCAPE’s similarity metric can then be used as an edge-length in network analyses of cluster similarity. We evaluated networks using an edge-length cutoff from 0.1-0.9 with a step of 0.1 (Fig. S4). We found networks with an edge-length cutoff of 0.4-0.6 to be similar and based further analyses on a cutoff of 0.5. For BGCs of interest, we supplemented BiG-SCAPE’s approach to identifying homologous BGCs with visualize inspection of microsyteny and blast-based analyses using NCBI’s BLAST+, v2.3.0 (Camacho et al., 2009). Similar sequences in microsynteny analyses were defined as at least 100 bp in length, at least 30 percent similarity, and an expectation value threshold of 0.01.

### Identification and characterization of secondary metabolite production General experimental procedures

The ^1^H NMR data were collected using a JOEL ECS-400 spectrometer, which was equipped with a JOEL normal geometry broadband Royal probe, and a 24-slot autosampler, and operated at 400 MHz. HRESIMS experiments utilized either a Thermo LTQ Orbitrap XL mass spectrometer or a Thermo Q Exactive Plus (Thermo Fisher Scientific); both were equipped with an electrospray ionization source. A Waters Acquity UPLC (Waters Corp.) was utilized for both mass spectrometers, using a BEH C_18_ column (1.7 μm; 50 mm x 2.1 mm) set to a temperature of 40oC and a flow rate of 0.3 ml/min. The mobile phase consisted of a linear gradient of CH_3_CN-H_2_O (both acidified with 0.1% formic acid), starting at 15% CH_3_CN and increasing linearly to 100% CH_3_CN over 8 min, with a 1.5 min hold before returning to the starting condition. The HPLC separations were performed with Atlantis T3 C_18_ semi-preparative (5 μm; 10 x 250 mm) and preparative (5 μm; 19 x 250 mm) columns, at a flow rate of 4.6 ml/min and 16.9 ml/min, respectively, with a Varian Prostar HPLC system equipped with a Prostar 210 pumps and a Prostar 335 photodiode array detector (PDA), with the collection and analysis of data using Galaxie Chromatography Workstation software. Flash chromatography was performed on a Teledyne ISCO Combiflash Rf 200 and monitored by both ELSD and PDA detectors.

### Chemical characterization

To identify the secondary metabolites that were biosynthesized by *A. fumigatus, A. oerlinghausenensis*, and *A. fischeri*, these strains were grown as large-scale fermentations to isolate and characterize the secondary metabolites. To inoculate oatmeal cereal media (Old fashioned breakfast Quaker oats), agar plugs from fungal stains grown on potato dextrose agar; difco (PDA) were excised from the edge of the Petri dish culture and transferred to separate liquid seed media that contained 10 ml YESD broth (2% soy peptone, 2% dextrose, and 1% yeast extract; 5 g of yeast extract, 10 g of soy peptone, and 10 g of D-glucose in 500 ml of deionized H_2_O) and allowed to grow at 23°C with agitation at 100 rpm for three days. The YESD seed cultures of the fungi were subsequently used to inoculate solid-state oatmeal fermentation cultures, which were either grown at room temperature (approximately 23°C under 12h light/dark cycles for 14 days), 30°C, or 37°C; all growths at the latter two temperatures were carried out in an incubator (VWR International) in the dark over four days. The oatmeal cultures were prepared in 250 ml Erlenmeyer flasks that contained 10 g of autoclaved oatmeal (10 g of oatmeal with 17 ml of deionized H_2_O and sterilized for 15–20 minutes at 121°C). For all fungal strains three flasks of oatmeal cultures were grown at all three temperatures, except for *A. oerlinghausenensis* (CBS 139183^T^) at room temperature and *A. fumigatus* (Af293) at 37°C. For CBS 139183^T^, the fungal cultures were grown in four flasks, while for Af293 eight flasks were grown in total. The growths of these two strains were performed differently from the rest because larger amounts of extract were required in order to perform detailed chemical characterization. The cultures were extracted by adding 60 ml of (1:1) MeOH-CHCl_3_ to each 250 ml flask, chopping thoroughly with a spatula, and shaking overnight (∼ 16 h) at ∼ 100 rpm at room temperature. The culture was filtered *in vacuo*, and 90 ml CHCl_3_ and 150 ml H_2_O were added to the filtrate. The mixture was stirred for 30 min and then transferred to a separatory funnel. The organic layer (CHCl_3_) was drawn off and evaporated to dryness *in vacuo*. The dried organic layer was reconstituted in 100 ml of (1:1) MeOH–CH_3_CN and 100 ml of hexanes, transferred to a separatory funnel, and shaken vigorously. The defatted organic layer (MeOH–CH_3_CN) was evaporated to dryness *in vacuo*.

To isolate compounds, the defatted extract was dissolved in CHCl_3_, absorbed onto Celite 545 (Acros Organics), and fractioned by normal phase flash chromatography using a gradient of hexane-CHCl_3_-MeOH. *Aspergillus fischeri* strain NRRL 181 was chemically characterized previously (Knowles et al., 2019; Mead et al., 2019a). *A. fumigatus* strain Af293, grown at 37°C, was subjected to a 12g column at a flow rate of 30 ml/min and 61.0 column volumes, which yielded four fractions. Fraction 2 was further purified via preparative HPLC using a gradient system of 30:70 to 100:0 of CH_3_CN-H_2_O with 0.1% formic acid over 40 min at a flow rate of 16.9 ml/min to yield six subfractions. Subfractions 1, 2 and 5, yielded cyclo(L-Pro-L-Leu) (**1**) (Li et al., 2008) (0.89 mg), cyclo(L-Pro-L-Phe) (**2**) (Campbell et al., 2009) (0.71 mg), and monomethylsulochrin (**3**) (Ma et al., 2004) (2.04 mg), which eluted at approximately 5.7, 6.3, and 10.7 min, respectively. Fraction 3 was further purified via preparative HPLC using a gradient system of 40:60 to 65:35 of CH_3_CN-H_2_O with 0.1% formic acid over 30 min at a flow rate of 16.9 ml/min to yield four subfractions. Subfractions 1 and 2 yielded pseurotin A (**4**) (Wang et al., 2011) (12.50 mg) and bisdethiobis(methylthio)gliotoxin (**5**) (Afiyatullov et al., 2005) (13.99 mg), which eluted at approximately 7.5 and 8.0 min, respectively.

*A. fumigatus* strain CEA10, grown at 37°C, was subjected to a 4g column at a flow rate of 18 ml/min and 90.0 column volumes, which yielded five fractions. Fraction 1 was purified via preparative HPLC using a gradient system of 50:50 to 100:0 of CH_3_CN-H_2_O with 0.1% formic acid over 45 min at a flow rate of 16.9 ml/min to yield eight subfractions. Subfraction 1, yielded fumagillin (**6**) (Halász et al., 2000) (1.69 mg), which eluted at approximately 18.5 min. Fraction 2 was purified via semi-preparative HPLC using a gradient system of 35:65 to 80:20 of CH_3_CN-H_2_O with 0.1% formic acid over 30 min at a flow rate of 4.6 ml/min to yield 10 subfractions. Subfraction 5 yielded fumitremorgin C (**7**) (Kato et al., 2009) (0.25 mg), which eluted at approximately 15.5 min. Fraction 3 was purified via preparative HPLC using a gradient system of 40:60 to 100:0 of CH_3_CN-H_2_O with 0.1% formic acid over 30 min at a flow rate of 16.9 ml/min to yield nine subfractions. Subfraction 2 yielded pseurotin A (**4**) (1.64 mg), which eluted at approximately 7.3 min.

*Aspergillus oerlinghausenensis* strain CBS 139183^T^, grown at RT, was subjected to a 4g column at a flow rate of 18 ml/min and 90 column volumes, which yielded 4 fractions. Fraction 3 was further purified via preparative HPLC using a gradient system of 35:65 to 70:30 of CH_3_CN-H_2_O with 0.1% formic acid over 40 min at a flow rate of 16.9 ml/min to yield 11 subfractions. Subfractions 3 and 10 yielded spiro [5H,10H-dipyrrolo[1,2-a:1′,2′-d]pyrazine-2-(3H),2′-[2H]indole]-3′,5,10(1′H)-trione (**8**) (Wang et al., 2008) (0.64 mg) and helvolic acid (**9**) (Zhao et al., 2010) (1.03 mg), which eluted at approximately 11.5 and 39.3 min, respectively. (see NMR supporting information; figshare: 10.6084/m9.figshare.12055503).

### Metabolite profiling by mass spectrometry

The metabolite profiling by mass spectrometry, also known as dereplication, was performed as stated previously (El-Elimat et al., 2013). Briefly, ultraperformance liquid chromatography-photodiode array-electrospray ionization high resolution tandem mass spectrometry (UPLC-PDA-HRMS-MS/MS) was utilized to monitor for secondary metabolites across all strains (Af293, CEA10, CEA17, CBS 139183^T^, NRRL 181, NRRL 4161, and NRRL 4585). Utilizing positive-ionization mode, ACD MS Manager with add-in software IntelliXtract (Advanced Chemistry Development, Inc.; Toronto, Canada) was used for the primary analysis of the UPLC-MS chromatograms. The data from 19 secondary metabolites are provided in the Supporting Information (see Dereplication table; figshare: 10.6084/m9.figshare.12055503), which for each secondary metabolite lists: molecular formula, retention time, UV-absorption maxima, high-resolution full-scan mass spectra, and MS-MS data (top 10 most intense peaks).

### Metabolomics analyses

Principal component analysis (PCA) analysis was performed on the UPLC-MS data. Untargeted UPLC-MS datasets for each sample were individually aligned, filtered, and analyzed using MZmine 2.20 software (https://sourceforge.net/projects/mzmine/) (Pluskal et al., 2010). Peak detection was achieved using the following parameters, *A. fumigatus* at (Af293, CEA10, and CEA17): noise level (absolute value), 1×10^6^; minimum peak duration, 0.05 min; *m/z* variation tolerance, 0.05; and *m/z* intensity variation, 20%; *A. fischeri* (NRRL 181, NRRL 4161, and NRRL 4585): noise level (absolute value), 1×10^6^; minimum peak duration, 0.05 min; *m/z* variation tolerance, 0.05; and *m/z* intensity variation, 20%; and all strains (Af293, CEA10, CEA17, CBS 139183^T^, NRRL 181, NRRL 4161, and NRRL 4585): noise level (absolute value), 7×10^5^; minimum peak duration, 0.05 min; *m/z* variation tolerance, 0.05; and *m/z* intensity variation, 20%. Peak list filtering and retention time alignment algorithms were used to refine peak detection. The join algorithm integrated all sample profiles into a data matrix using the following parameters: *m/z* and retention time balance set at 10.0 each, *m/z* tolerance set at 0.001, and RT tolerance set at 0.5 mins. The resulting data matrix was exported to Excel (Microsoft) for analysis as a set of *m/z* – retention time pairs with individual peak areas detected in triplicate analyses. Samples that did not possess detectable quantities of a given marker ion were assigned a peak area of zero to maintain the same number of variables for all sample sets. Ions that did not elute between 2 and 8 minutes and/or had an *m/z* ratio less than 200 or greater than 800 Da were removed from analysis. Relative standard deviation was used to understand the quantity of variance between the technical replicate injections, which may differ slightly based on instrument variance. A cutoff of 1.0 was used at any given *m/z* – retention time pair across the technical replicate injections of one biological replicate, and if the variance was greater than the cutoff, it was assigned a peak area of zero. Final chemometric analysis, data filtering (Caesar et al., 2018) and PCA was conducted using Sirius, v10.0 (Pattern Recognition Systems AS) (Kvalheim et al., 2011), and dendrograms were created with Python. The PCA scores plots were generated using data from either the three individual biological replicates or the averaged biological replicates of the fermentations. Each biological replicate was plotted using averaged peak areas obtained across four replicate injections (technical replicates).

## Data Availability

Sequence reads and associated genome assemblies generated in this project are available in NCBI’s GenBank database under the BioProject PRJNA577646. Additional descriptions of the genomes including predicted gene boundaries will become available through the Figshare repository 10.6084/m9.figshare.12055503 upon publication. The Figshare repository is also populated with other data generated from genomic and natural products analysis. Among genomic analyses, we provide information about predicted BGCs, results associated with network-based clustering of BGCs into cluster families, phylogenomic data matrices, and others. Among natural products analysis, we provide information that supports methods, and results, including NMR spectra.

## Funding

JLS and AR are supported by the Howard Hughes Medical Institute through the James H. Gilliam Fellowships for Advanced Study program. AR has additional support from a Discovery Grant from Vanderbilt University. GHG is supported by the Brazilian funding agencies Fundacão de Amparo a Pesquisa do Estado de São Paulo (FAPESP 2016/07870-9) and Conselho Nacional de Desenvolvimento Cientifico e Tecnologico (CNPq). NHO is supported by the National Cancer Institute (P01 CA125066). SLK and CDR were supported in part by the National Institutes of Health via the National Center for Complementary and Integrative Health (F31 AT010558) and the National Institute of General Medical Sciences (T34 GM113860), respectively.

## Acknowledgements

We thank the labs of Rokas, Oberlies, and Goldman for helpful discussion and support of this work.

## Supplementary legends

**Fig. S1. Metrics of genomes assembly quality and number of predicted gene.** Genome size, N50, number of genes, number of scaffolds and percent single copy BUSCO genes (scBUSCO) are depicted here. Examination of metrics reveal genomes are of sufficient quality for comparative genomics purposes. Of concern, we noted A. oerlinghausenensis CBS 139183^T^ was assembled into 5,300 contigs; however, we evaluated the assembly’s N50 value and gene content completeness (461,327 base pairs and 98.90% BUSCO genes present in single copy, respectively) and found the *A. oerlinghausenensis* genome is suitable for comparative genomic analyses.

**Fig. S2. A reconstructed evolutionary history and timetree of *A. fumigatus* and its closest relatives.** (top) Divergence times were estimated using a concatenated matrix of 3,041 genes (5,602,272 sites). Blue bars at each node correspond to the 95% divergence time confidence interval. Divergence times and confidence intervals for each internode are shown on the right side of the figure. (bottom) A phylogeny where branch lengths represent substitutions per site rather than geologic time. A cladogram is drawn to the right of the phylogeny to clarify divergences where branch lengths are short (e.g., among strains of *A. fischeri*).

**Fig. S3. Moderate edge cut-off lengths resulted in qualitatively similar networks.** Networks using edge cut-offs ranging from 0.1-0.9 with a step of 0.1 were evaluated. Networks from 0.4 to 0.6 were qualitatively similar. Thus, we used cluster families inferred from the network using an edge cut-off of 0.5.

**Fig. S4. Species and strain heterogeneity among BGC cluster presence and absence.** (A) Examination of the number of strains in each cluster family reveal a wide variation. For example, all 10 genomes are represented in 17 cluster families; in contrast, 18 cluster families have only one BGC. (B) Species occupancy among cluster families reveal *A. fischeri* has the largest number of unique BGCs followed by *A. fumigatus* and *A. oerlinghausenensis*. (C) Strain-level presence and absence patterns among cluster families reveal substantial species and strain heterogeneity. Cluster family identifiers are shown along the x-axis. Genus and species names are written using the following abbreviations: *Afum*: *A. fumigatus*; *Aoer*: *A. oerlinghausenensis*; *Afis*: *A. fischeri*.

**Fig. S5. Gene family presence and absence follows a similar pattern to BGCs.** Orthogroups were used as proxies for gene families. A strain-level UpSet plot for gene family presence and absence patterns reveals heterogeneity among strains.

**Fig. S6. Structures of isolated fungal metabolites.** Secondary metabolites produced in sufficient quantity were isolated for structural determination. The structures of compounds are correlated to each strain from where they were isolated.

**Fig. S7. Principal component analysis of BGC presence and absence and secondary metabolite profiles mirror one another.** (A) Principal component analysis of BGC presence and absence reveal that each species is distinct from the other. Furthermore, *A. oerlinghausenensis* is between *A. fischeri* and *A. fumigatus*. (B, C) Broadly, similar patterns of species relationships in principal component space are observed among all metabolites produced at 37°C and 30°C. (D, E) Similar results were observed for isolatable metabolites.

**Fig. S8. Small sequence divergences in the trypacidin BGC are associated with the production of trypacidin or the lack thereof.** The trypacidin BGC is found in *A. fumigatus* strains Af293 and CEA10, *A. oerlinghausenensis* CBS 139183^T^, and *A. fischeri* NRRL 181. Evidence of trypacidin biosynthesis is found in all isolates with the exception of *A. fischeri* NRRL 181. The absence of trypacidin biosynthesis is associated with the absence of a RING finger gene in *A. fischeri* NRRL 181. Black and white squares correspond to the presence or absence of the associated secondary metabolite, respectively. Genus and species names are written using the following abbreviations: *Afum*: *A. fumigatus*; *Aoer*: *A. oerlinghausenensis*; *Afis*: *A. fischeri*.

## Notes

### Competing Interest Statement

The authors have declared no competing interest.

