## Supplementary Figures for "Biosynthetic gene clusters, secondary metabolite profiles, and cards of virulence in the closest nonpathogenic relatives of *Aspergillus fumigatus*"

Supplementary figure 1

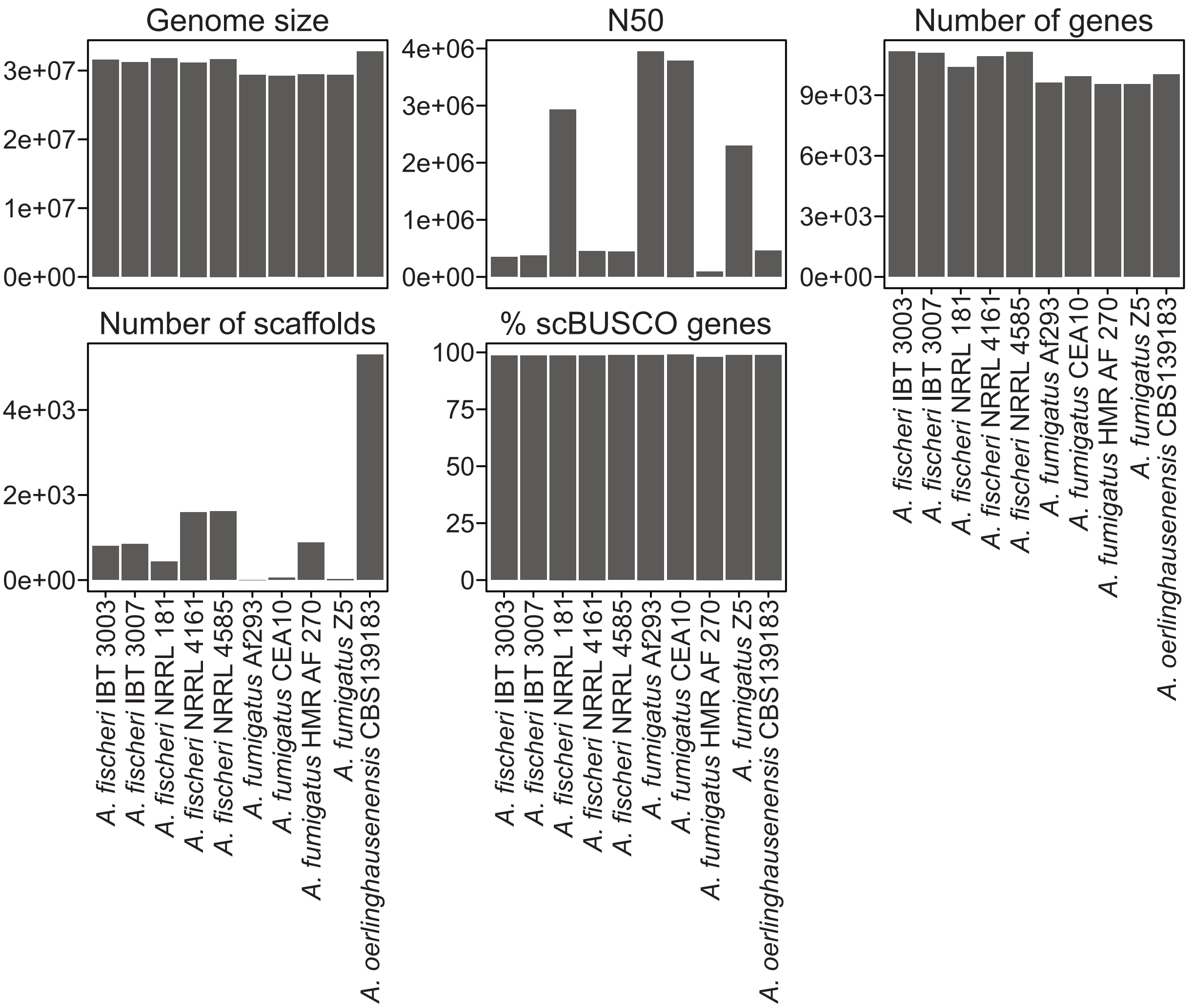

Supplementary figure 2

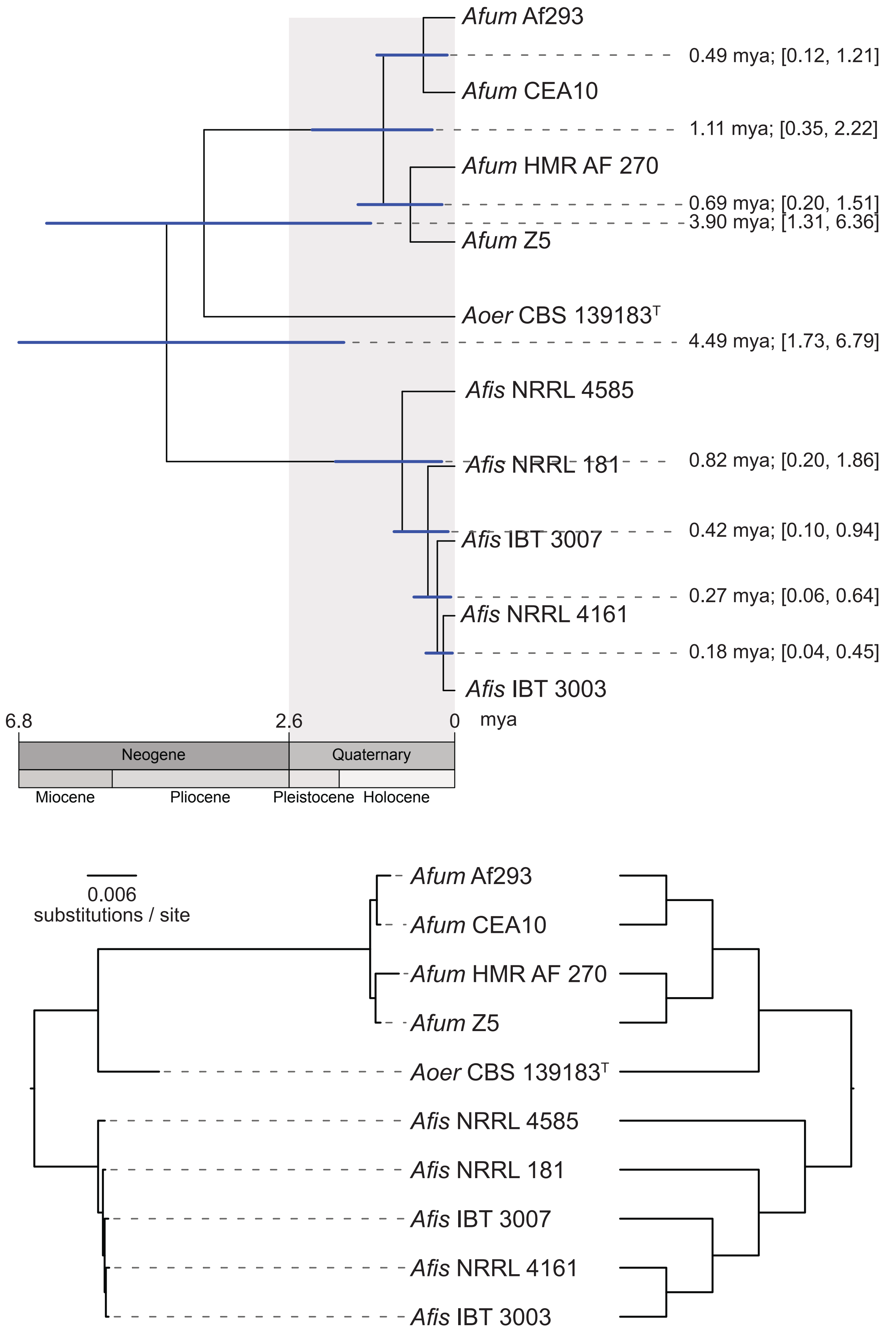

Supplementary figure 3

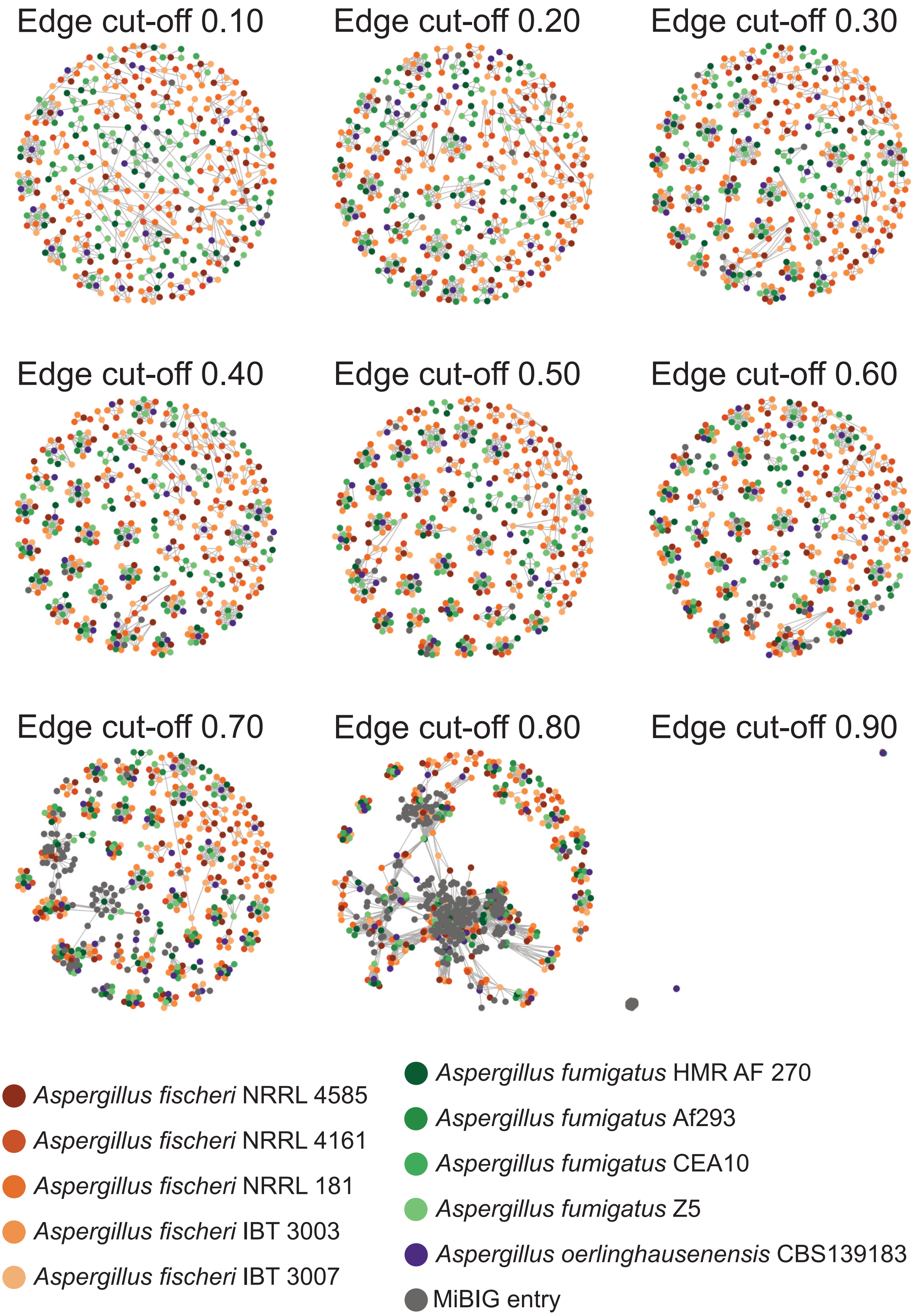

Supplementary figure 4

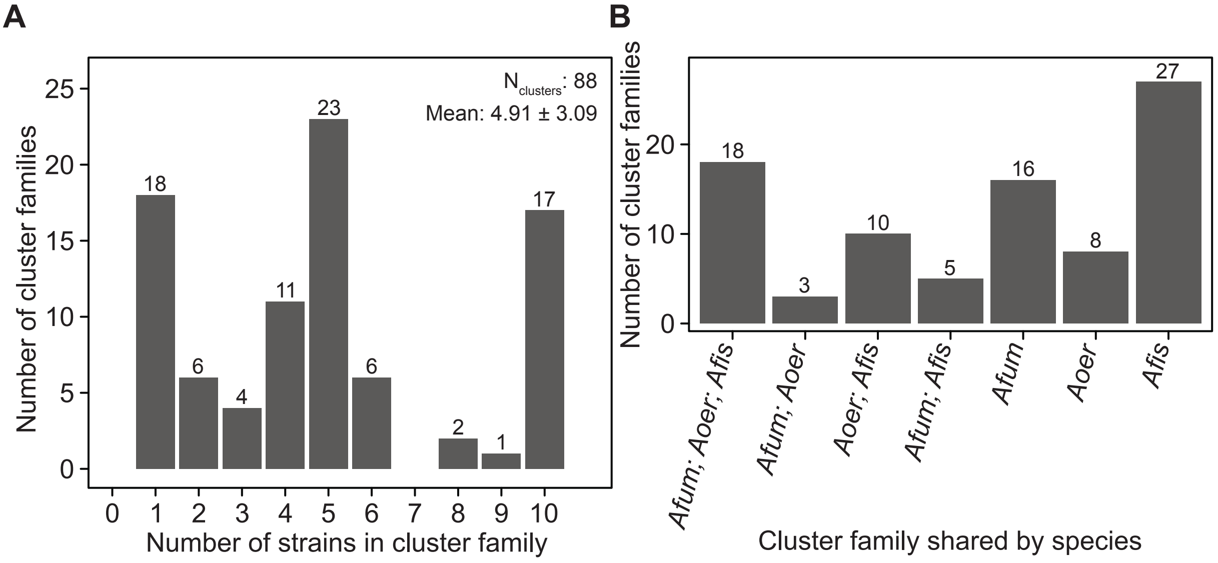

Supplementary figure 5

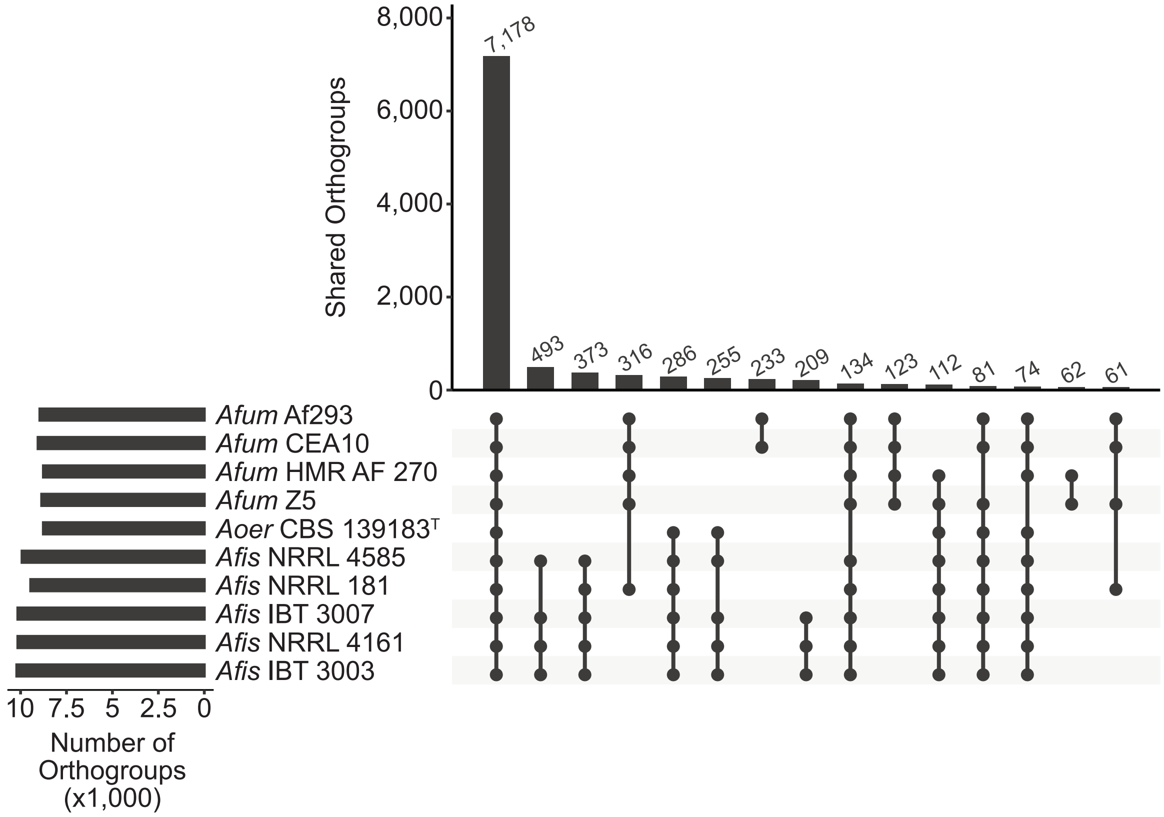

Supplementary figure 6

### *A. fumigatus* Af293

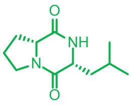

Cyclo (L-Pro-L-Leu) **(1)**

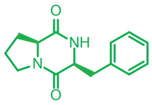

Cyclo (L-Pro-L-Phe) **(2)**

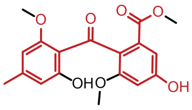

Monomethylsulochrin **(3)**

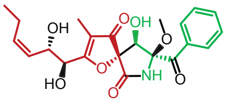

Pseurotin A **(4)**

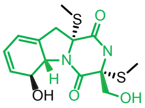

Bisdethiobis(methylthio)gliotoxin **(5)**

### *A. fumigatus* CEA10

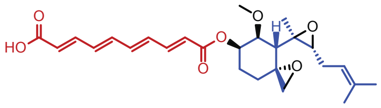

Fumagillin **(6)**

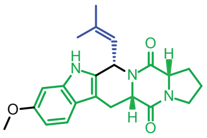

Fumitremorgin C **(7)**

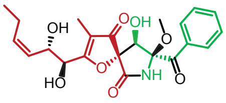

Pseurotin A **(4)**

### *A. oerlinghausenensis* CBS 139183[T]

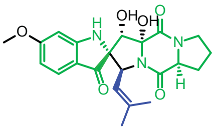

spiro [5H,10H-dipyrrolo[1,2-a:1',2'-d]pyrazine-2-(3H),2'-[2H]indole]-3',5,10(1'H)-trione **(8)**

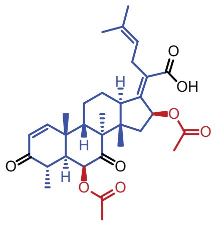

Helvolic acid **(9)**

### *A. fischeri* strains NRRL 181, NRRL 4585, NRRL 4161

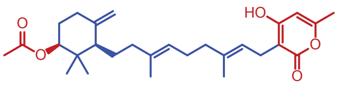

Sartorypyrone A **(10)**

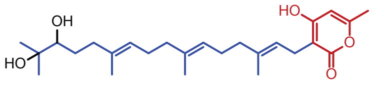

Sartorypyrone E **(11)**

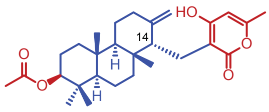

14-epi-aszonapyrone A **(12)**

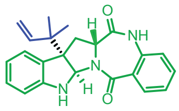

Aszonalenin **(13)**

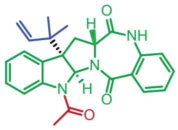

Acetylaszonalenin **(14)**

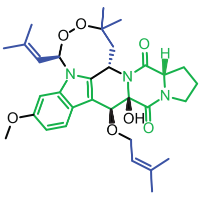

Fumitremorgin A **(15)**

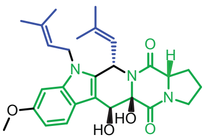

Fumitremorgin B **(16)**

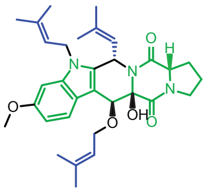

13-*O*-prenyl-fumitremorgin B **(17)**

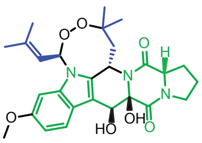

Verruculogen **(18)**

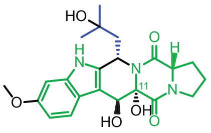

11-epimer of verrucologen TR2 **(19)**

**Color Key**

- Terpene
- Polyketide Synthase
- Non-ribosomal Peptide Synthetase

Supplementary figure 7

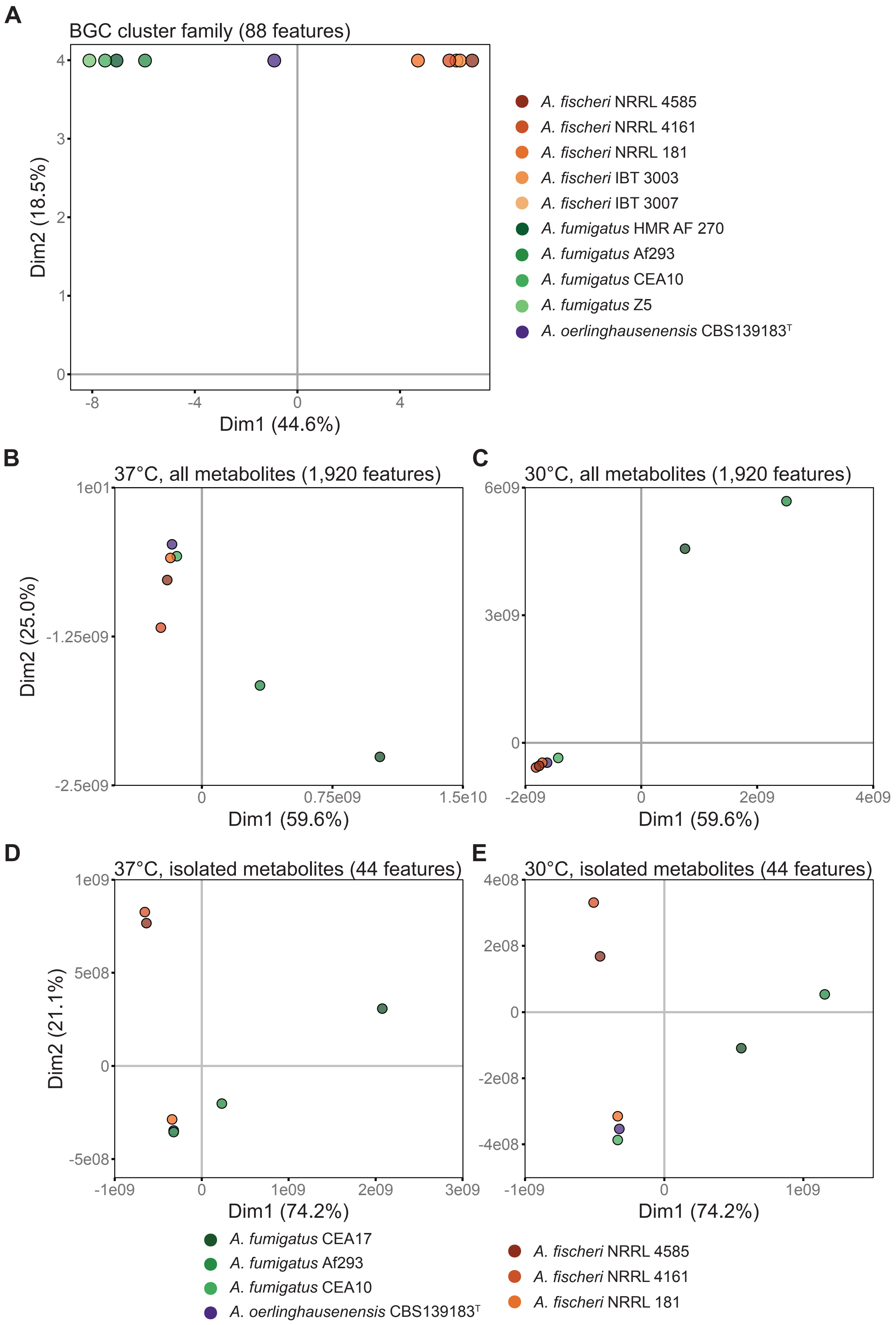

Supplementary figure 8

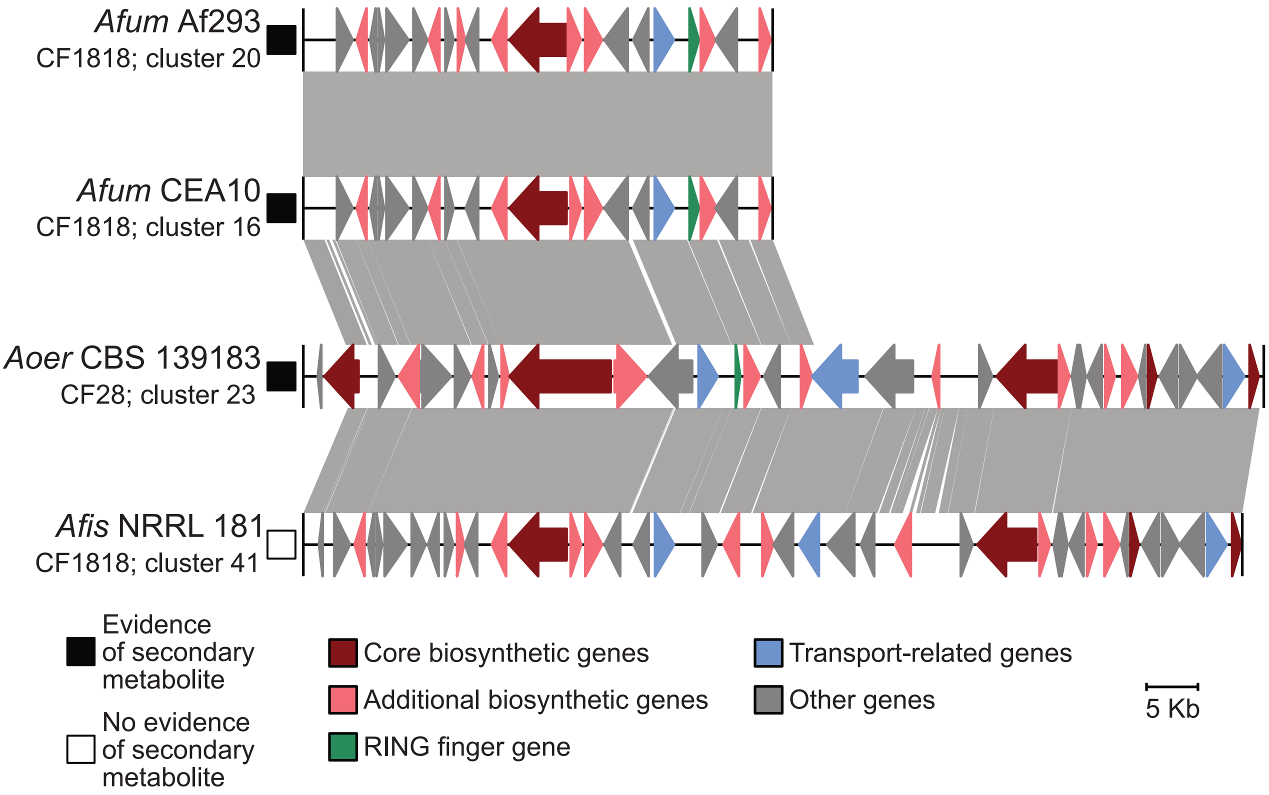
